# Fasting reverses PLN R14del-mediated cardiomyopathy through lysosomal reactivation

**DOI:** 10.64898/2026.03.24.713684

**Authors:** Iris Gooijers, Aleksandra Arning, Cecilia de Heus, Ursula Heins-Marroquin, Phong D. Nguyen, Hessel Honkoop, Tim Verhagen, Mostafa M. Mokhles, Anneline S. J. M. te Riele, Magdalena Harakalova, Gijs van Haaften, Linda W. van Laake, Lukas C. Kapitein, Nalan Liv, Jeroen Bakkers

**Author notes:** Senior author and lead contact (Jeroen Bakkers).

## Abstract

Genetic cardiomyopathies consist of a heterogeneous group of myocardial disorders caused by variants that disrupt key regulators of cardiac structure and function. Variants in *PLN,* encoding phospholamban (PLN), the main inhibitor of the sarco/endoplasmic reticulum Ca²⁺-ATPase 2a (SERCA2a), have been linked to both dilated cardiomyopathy (DCM) and arrhythmogenic cardiomyopathy (ACM). Among these, the PLN Arg14del (R14del) variant is the most prevalent. PLN R14del cardiomyopathy is characterized by the accumulation of large perinuclear PLN aggregates in cardiomyocytes of end-stage heart failure tissue. However, the mechanisms driving PLN aggregate formation and their role in disease progression remain unresolved.

Using a humanized *plna* R14del zebrafish model, left ventricular tissue from end-stage PLN R14del cardiomyopathy patients and pharmacological modeling in wild type (WT) cardiac slices, we demonstrate that previously described PLN aggregates represent accumulated sarcoplasmic reticulum (SR)-derived PLN-containing vesicles that form due to impaired SERCA2a activity and increased cytosolic Ca²⁺ levels. Furthermore, these SR-derived vesicles often localize adjacent to lysosomes. Interestingly, Ca^2+^ dysregulation in *plna* R14del hearts leads to reduced lysosomal function, resulting in SR-derived vesicle accumulation at the microtubule organizing center (MTOC). This perinuclear accumulation induces microtubule aster formation and subsequent cellular disorganization, including sarcomere misalignment and nuclear deformation. Strikingly, reactivation of lysosomal function through fasting reduces SR-derived vesicle accumulation, restores microtubule integrity, and rescues cellular organization in *plna* R14del zebrafish hearts.

Together, these findings identify impaired lysosomal clearance of SR-derived vesicles and the resulting microtubule disorganization as key pathological mechanisms driving PLN R14del cardiomyopathy. Additionally, our results highlight lysosomal reactivation as a promising potential therapeutic strategy to halt or reverse PLN R14del cardiomyopathy progression.

**Main findings:**

- PLN aggregates in PLN R14del cardiomyopathy represent SR-derived vesicles formed due to Ca²⁺ dysregulation.
- These SR-derived vesicles often localize perinuclearly at the microtubule organizing center (MTOC), where they are positioned adjacent to lysosomes.
- Ca^2+^ dysregulation leads to lysosomal dysfunction which drives vesicle accumulation responsible for microtubule remodeling and pathological cellular rearrangements.
- Lysosomal reactivation restores vesicle clearance and rescues cardiomyocyte structure.

## Introduction

Cardiomyopathies consist of a heterogeneous group of myocardial diseases that largely contribute to cardiovascular morbidity and mortality worldwide. To date, multiple cardiomyopathy subtypes have been identified, among which dilated cardiomyopathy (DCM, prevalence ∼1:250), hypertrophic cardiomyopathy (HCM, prevalence ∼1:500), and arrhythmogenic cardiomyopathy (ACM, prevalence ∼1:1250) are the most common (Abbas et al., 2024; Arbelo et al., 2023; Austin et al., 2019; E. McNally et al., 1993; E. M. McNally & Mestroni, 2017). Patients are at risk for heart failure, ventricular arrhythmias, and sudden cardiac death. Approximately half of all reported cardiomyopathy cases are caused by inherited genetic variants (Abbas et al., 2024; Arbelo et al., 2023; Austin et al., 2019; E. McNally et al., 1993; E. M. McNally & Mestroni, 2017). These variants are found in genes encoding proteins critical for desmosomes, adherens junctions, cytoskeletal structure, and ion channels (Austin et al., 2019; Gerull & Brodehl, 2020). Variants in *PLN,* encoding phospholamban (PLN), a key regulator in Ca^2+^ handling (Lazzarini et al., 2015; van der Zwaag et al., 2012), are linked to DCM and ACM. PLN protein localizes to the sarcoplasmic reticulum (SR) and nuclear envelope (NE), where it regulates Ca²⁺ homeostasis by inhibiting sarco/endoplasmic reticulum Ca²⁺-ATPase 2a (SERCA2a), the key enzyme responsible for Ca²⁺ reuptake during excitation-contraction coupling (Berridge, 2003; Bers, 2002; Gilbert et al., 2020; A. Z. Wu et al., 2016).

Over the past three decades, several disease causing variants in the *PLN* gene have been identified, among which the deletion of arginine 14 (R14del) is the most prevalent (Hof et al., 2019; Schmitt et al., 2003). To date, only heterozygous carriers of the PLN R14del variant have been identified. Clinically, despite underlying mechanisms remaining poorly understood, patients carrying the PLN R14del variant may present with a DCM-like phenotype, characterized by ventricular dilation and systolic dysfunction, or an ACM-like phenotype marked by ventricular arrhythmias and fibrofatty replacement (te Rijdt et al., 2016). Remarkably, these phenotypic traits often overlap in a single patient (te Rijdt et al., 2016). On a molecular level, PLN R14del has been widely described as a super-inhibitor of SERCA2a, causing impaired Ca^2+^ reuptake in the SR and elevated cytosolic Ca^2+^ levels (Haghighi et al., 2006, 2021; Kumar et al., 2023; Vafiadaki et al., 2022). However, conflictingly, other studies show accelerated SERCA2a activity (Stege et al., 2024; Vafiadaki et al., 2023). Despite these discrepancies, all models consistently demonstrate abnormal intracellular Ca²⁺ handling. In addition to Ca²⁺ dysregulation, PLN R14del cardiomyopathy is characterized by perinuclear PLN aggregates (Eijgenraam et al., 2021; te Rijdt et al., 2016, 2017; Vafiadaki et al., 2024), which were recently defined as aberrantly clustered SR membranes (Stege et al., 2023). Despite their frequent occurrence in various PLN R14del disease models, the mechanisms driving the formation of these structures and their contribution to disease progression remain unresolved. This limits mechanistic understanding and the development of targeted therapies for PLN R14del cardiomyopathy. Specifically, it is unclear whether these structures are a byproduct of chronic Ca²⁺ dysregulation and/or whether they contribute directly to cellular dysfunction and cardiac remodeling involved in PLN R14del cardiomyopathy disease progression.

To gain further insight into PLN R14del disease progression, we previously generated a humanized Pln R14del zebrafish model (*plna* R14del/R14del; *plnb* +/+) (Kamel et al., 2021; Tessadori et al., 2018), reflecting the heterozygous PLN R14del variant identified in human patients. Hereafter we refer to this model as *plna* R14del zebrafish. Our *plna* R14del zebrafish model recapitulates key hallmarks of the disease in humans, including SERCA2a inhibition, progressive cardiac dysfunction, and fibrofatty replacement characteristic of end-stage cardiomyopathy in two years old zebrafish. In the present study, we leverage our humanized *plna* R14del zebrafish model, together with cardiac tissue obtained from patients with end-stage cardiomyopathy carrying the PLN R14del variant and pharmacological modeling in wild type (WT) cardiac slices, to investigate the mechanisms underlying perinuclear SR clustering and its downstream consequences. We show that the aberrant PLN^+^ perinuclear SR structures originate from SR-derived vesicles formed in response to impaired SERCA2a activity and elevated cytosolic Ca²⁺. These vesicles are often localized adjacent to lysosomes. Chronic Ca²⁺ dysregulation in *plna* R14del hearts leads to defective lysosomal clearance, resulting in progressive accumulation of SR-derived vesicles at the microtubule organizing center (MTOC), accompanied by microtubule remodeling, sarcomere disorganization, and nuclear deformation. Importantly, restoring lysosomal function through fasting reduces vesicle accumulation and rescues cardiomyocyte structure. Collectively, these findings reveal impaired vesicle clearance by lysosomes as a key driver of PLN R14del cardiomyopathy progression and identify lysosomal reactivation as a potential therapeutic strategy.

## Results

### Ca^2+^ dysregulation in PLN R14del cardiomyopathy drives progressive SR-derived vesicle accumulation

A hallmark of PLN R14del cardiomyopathy is the accumulation of perinuclear PLN aggregates (Eijgenraam et al., 2021; te Rijdt et al., 2016, 2017; Vafiadaki et al., 2024). These PLN aggregates have recently been identified as aberrantly clustered SR membranes in both mouse and human PLN R14del cardiomyocytes (Stege et al., 2023). To further characterize these aberrant structures, we used left ventricular (LV) tissue obtained from patients with end-stage cardiomyopathy carrying the PLN R14del variant. Histological examination of these samples demonstrated extensive fibrofatty replacement, as confirmed by Picrosirius red staining, consistent with end-stage disease (**Figure 1A**). High-resolution immunofluorescence imaging revealed that PLN occasionally localized in vesicle-like structures near the nucleus (**Figure 1B**). Further examination of tissue from the same patient showed that similar vesicle-like structures are also incorporated within larger perinuclear PLN clusters (**Figure 1C**). Together, this suggests that the structures previously described as PLN aggregates result from the accumulation of SR-derived vesicles, which eventually give rise to larger aberrant membranous PLN clusters, thereby providing a mechanistic basis for the aberrantly clustered SR membranes reported previously (Stege et al., 2023).

**Figure 1.**
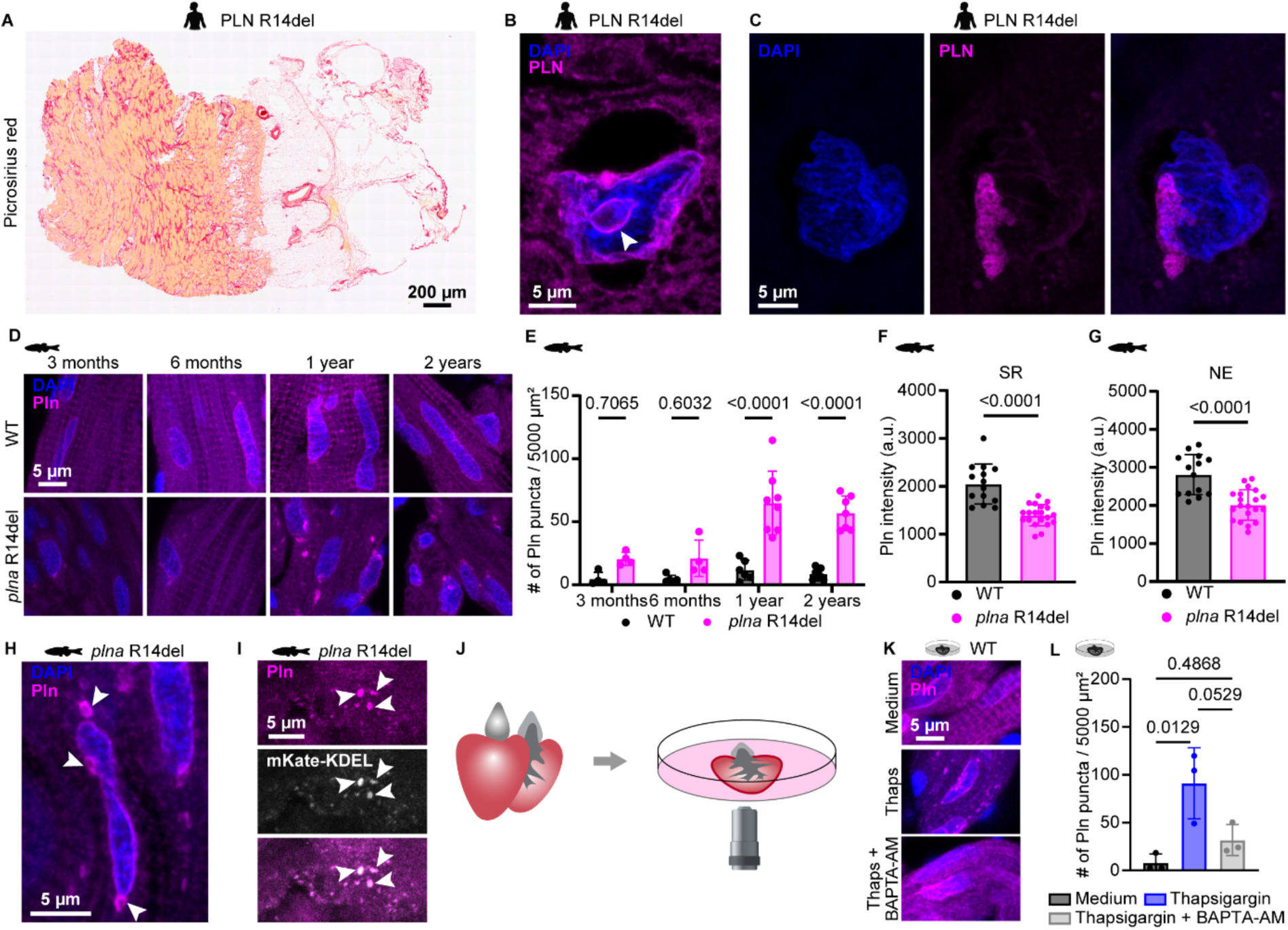
Ca^2+^ dysregulation in PLN R14del cardiomyopathy drives progressive SR-derived vesicle accumulation in zebrafish and human cardiomyocytes. **A.** Representative image of a Picrosirius red staining in left ventricle (LV) from patients with end-stage cardiomyopathy carrying the PLN R14del variant. Data shown are representative for n = 4 individuals. Scale bar = 200 μm. **B-C.** Representative images of immunofluorescent staining against DAPI and PLN in left ventricle (LV) from patients with end-stage cardiomyopathy carrying the PLN R14del variant showing **(B)** localization of PLN inside a vesicle (white arrowhead) and **(C)** perinuclear clustered PLN^+^ membrane-like structures (scale bars = 5 μm). Data shown are representative for n = 2 individuals. **D.** Representative images of immunofluorescent staining against DAPI and Pln on three months, six months, one year and two years old wild type (WT) and *plna* R14del zebrafish hearts. Data shown are representative for at least n = 3 individuals per group per timepoint. Scale bar = 5 μm. **E.** Quantification of Pln puncta per 5000 μm^2^. Data points represent averages of individual hearts. Data obtained from at least n = 3 individuals per group per timepoint. Error bars indicate mean ± s.d. Statistics were performed by two-way ANOVA. **F-G.** Quantification of Pln fluorescent intensity at the sarcoplasmic reticulum (SR) **(F)** and the nuclear envelop (NE) **(G)** in one year old WT and *plna* R14del zebrafish. Data points represent an individual measurement; multiple points originate from one individual (n = 6 WT and n = 8 *plna* R14del hearts). Error bars indicate mean ± s.d. Statistics were performed by two-tailed unpaired t-test. **H.** Representative images of immunofluorescent staining against DAPI and Pln in *plna* R14del zebrafish. White arrowheads indicate vesicles. Data shown are representative for n = 8 individuals. Scale bar = 5 μm. **I.** Representative images of immunofluorescent staining of Pln and mKate-KDEL in *plna* R14del Tg(*myl7*:mKate-KDEL;*exorh*:GFP) zebrafish. White arrowhead indicates a Pln and mKate-KDEL double positive punctas. Data shown are representative for n = 3 individuals. Scale bar = 5 μm. **J.** Schematic overview of the cardiac slice procedure; zebrafish hearts were extracted and sectioned into 150 µm thick slices using a vibratome. The slices were then cultured and subjected to specific drug treatments. **K.** Representative images of immunofluorescent staining against DAPI and Pln in WT zebrafish cardiac slices treated with medium, thapsigargin (100 μM) or thapsigargin (100 μM) + BAPTA-AM (100 μM). Data shown are representative for n = 3 individuals per group. Thaps = Thapsigargin; scale bar = 5 μm. **L.** Quantification of Pln puncta per 5000 μm^2^ after 5 hours of treatment. Data points represent average per individual (n = 3) per timepoint. Error bars indicate mean ± s.d. Statistics were performed by one-way ANOVA.

To further address when and how these PLN-containing vesicles appear we examined cardiac tissue from WT and *plna* R14del zebrafish at various ages. Zebrafish is a valid model to study PLN-induced cardiomyopathy as they recapitulate key characteristics of the end-stage disease, including SERCA2a inhibition, progressive cardiomyocytes dysfunction and fibrofatty replacement (Kamel et al., 2021). In WT cardiomyocytes, Pln consistently localized to the SR and NE at all examined timepoints (**Figure 1D**), which is in line with observations in other species (A. Z. Wu et al., 2016). Strikingly, we found that Pln puncta began to accumulate significantly in *plna* R14del cardiomyocytes from one year of age onwards, often localizing perinuclearly (**Figure 1D,E).** The accumulation of Pln in these puncta was accompanied by reduced Pln localization at the SR and NE (**Figure 1F,G**), recapitulating reduced PLN levels reported in cardiomyocytes of PLN R14del patients (Ragone et al., 2023). Higher magnifications showed the Pln puncta as rings, revealing these puncta as vesicles (**Figure 1H)**. This is in line with our observation of vesicle-like PLN localization near the nucleus in LV tissue obtained from patients with end-stage cardiomyopathy carrying the PLN R14del variant. To investigate the origin of these Pln^+^ vesicles, we generated a transgenic zebrafish line expressing a endoplasmic reticulum (ER)/SR-targeted mKate-KDEL fusion protein (Tg(*myl7*:mKate-KDEL;*exorh*:GFP)). The correct localization of this sensor was confirmed by evaluating co-localization between the mKate-KDEL sensor and Pln in WT zebrafish hearts (**Sup figure 1A**). As expected, we observed strong co-localization of Pln puncta with mKate-KDEL puncta in *plna* R14del zebrafish hearts, demonstrating that the observed Pln vesicles arise from the SR (**Figure 1I**).

Since *plna* R14del zebrafish cardiomyocytes have impaired Ca^2+^ reuptake into the SR as well as elevated intracellular Ca²⁺ levels (Kamel et al., 2021), we next asked whether Ca²⁺ dysregulation independent of the Pln mutation induces ER/SR stress which in turn could be responsible for the formation of SR-derived vesicles in *plna* R14del zebrafish. In line with this hypothesis, we observed that these Pln containing vesicles co-localized with Grp78 (**Sup figure 1B**), a marker for the unfolded protein response (UPR) which is activated upon ER/SR stress (Lee, 2005). To directly examine whether Ca²⁺ dysregulation drives the formation of Pln^+^ SR-derived vesicles, we used an *ex vivo* zebrafish cardiac slice system (Honkoop et al., 2021; Nguyen et al., 2023) and pharmacologically inhibited SERCA2a using thapsigargin (**Sup figure 1C**). Successful SERCA2a inhibition by thapsigargin was confirmed using cardiac slices expressing the Ca^2+^ sensor GCaMP6f (Nguyen et al., 2023; van Opbergen et al., 2018) (**Sup figure 1C-E**). Interestingly, thapsigargin treatment induced the accumulation of Pln puncta (**Figure 1J-L**), phenocopying PLN R14del-associated SR-derived vesicle accumulation. Notably, co-treatment with the Ca^2+^ chelator BAPTA-AM prevented Pln puncta accumulation, indicating that abnormally elevated cytoplasmic Ca^2+^ is necessary for this process (**Figure 1J-L)**. Together, our data demonstrate that perinuclear PLN clustering in PLN R14del hearts arises from SR-derived vesicle accumulation driven by Ca²⁺ dysregulation, rather than directly from the genetic variant itself.

### PLN R14del cardiomyopathy induced SR-derived vesicles localize adjacent to lysosomes

To investigate how PLN R14del and Ca^2+^ dysregulation contribute to SR-derived vesicle accumulation, we performed single-cell mRNA sequencing (scRNAseq) on cardiomyocytes isolated from one year old WT and *plna* R14del zebrafish hearts using the SOrting and Robot-assisted Transcriptome SEQuencing (SORT-seq) platform (Muraro et al., 2016). Uniform Manifold Approximation and Projection (UMAP) visualization revealed a main cardiomyocyte cluster, identified by high expression of cardiomyocyte marker genes, containing both WT and *plna* R14del cells (**Sup figure 2A-E**). *In silico* pseudobulk analysis of the SORT-seq data followed by gene ontology (GO) enrichment analysis revealed upregulation of genes involved in endocytosis, Golgi apparatus, and membrane coat formation in *plna* R14del hearts, pathways closely linked to endo-lysosomal biogenesis and function (**Figure 2A,B; Sup table 1-3**). Closer examination of individual genes confirmed upregulation of key lysosome-associated components, including core membrane and luminal lysosomal proteins (*lamp1b* & *man2b1*), factors involved in lysosomal acidification and membrane transport (*atp6v1e1b*, *slc36a1*, *slc9a6b* & *slc9a8*), and regulators of lysosomal signaling and trafficking (*lamtor5*, *rab33ba* & *sort1b*) (**Figure 2A,B; Sup table 1-3**). In contrast, genes involved in the proteasome complex were among the most strongly downregulated in *plna* R14del hearts (*psma4, psmb8a, psmb10, psmb13a, psmd4a* & *psme2*) (**Figure 2A,B; Sup table 1-3**). Additionally, no significant expression difference in autophagy pathway-related genes was observed (**Figure 2A,B; Sup table 1-3**).

**Figure 2.**
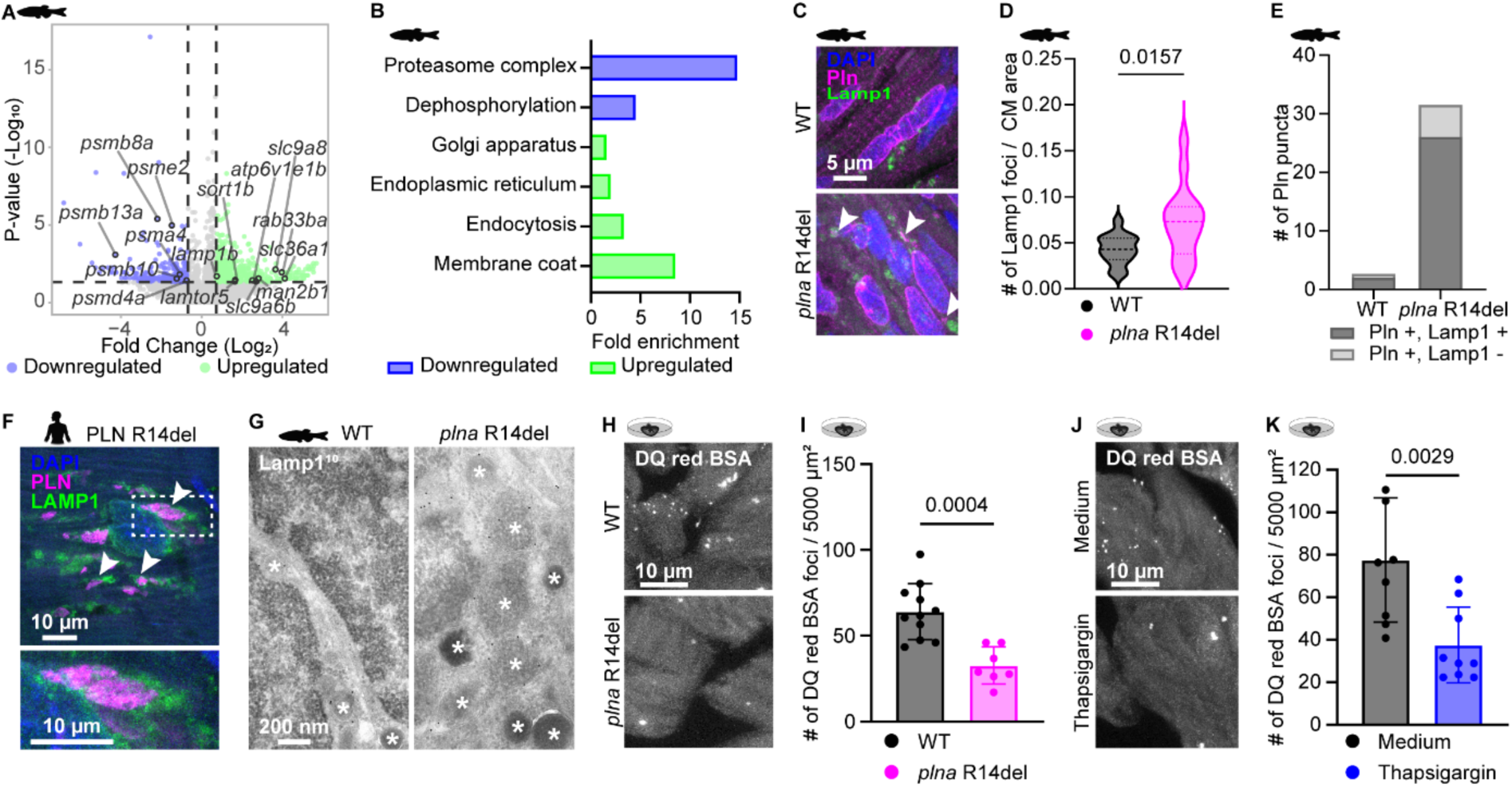
Ca^2+^ dysregulation impairs lysosomal function and promotes SR-derived vesicle accumulation. **A.** Volcano plot showing all significantly (p-value < 0.05) downregulated (n = 201) and upregulated (n = 433) genes in wild type (WT) and *plna* R14del zebrafish hearts with a log₂(fold change) higher than +/− 0.7. Each dot represents one gene; blue corresponds to downregulated genes and green to upregulated genes. The p-value cut off corresponds to 1.30 on the -log_10_ axis. **B.** Fold enrichment of downregulated and upregulated GO terms identified using the DAVID online GO analysis tool, when comparing WT and *plna* R14del zebrafish hearts in a pseudo-bulk dataset. **C.** Representative images of immunofluorescent staining against DAPI, Pln and Lamp1 in one year old WT and *plna* R14del zebrafish. Data shown are representative for n = 8 WT and n = 8 *plna* R14del zebrafish hearts. Scale bar = 5 μm. White arrowheads indicate Pln and Lamp1 co-localization. **D.** Quantification of number of Lamp1 foci divided by cardiomyocyte (CM) area. Statistics were performed by two-tailed unpaired t-test. Data obtained from n = 8 WT and n = 8 *plna* R14del zebrafish hearts. **E.** Quantification of Pln puncta co-localized with Lamp1 foci. Data obtained from n = 8 WT and n = 8 *plna* R14del zebrafish hearts. **F.** Representative image of immunofluorescent staining against DAPI, PLN and LAMP1 in left ventricle (LV) from patients with end-stage cardiomyopathy carrying the PLN R14del variant. Dashed box high lights the zoomed in area. Data shown are representative for n = 4 individuals. Scale bars = 10 μm. White arrowheads indicate PLN and LAMP1 co-localization. **G.** Representative images of immunoelectron microscopy against Lamp1 (10 nm gold) in one year old WT and *plna* R14del zebrafish hearts. * Indicate endo-lysosomal compartments. Data shown are representative for n = 3 individuals. Scale bar = 200 nm. **H.** Representative images of DQ red BSA BSA *ex vivo* assay in one year old WT and *plna* R14del hearts. Dots represent functional lysosomes. Data shown are representative for n = 4 WT and n = 3 *plna* R14del zebrafish hearts. Scale bar = 10 μm. **I.** Quantification of number of DQ red BSA foci per 5000 μm^2^. Data points represent an individual measurement per image; multiple points originate from one individual. Data obtained from n = 4 WT and n = 3 *plna* R14del zebrafish hearts. Error bars indicate mean ± s.d. Statistics were performed by two-tailed unpaired t-test. **J.** Representative images of DQ red BSA *ex vivo* assay in WT cardiac slices treated with medium or thapsigargin (100 μM). Dots represent functional lysosomes. Data shown are representative for n = 3 per group. Scale bar = 10 μm. **K.** Quantification of number of DQ red BSA foci per 5000 μm^2^. Data points represent an individual measurement per image; multiple points originate from one individual. Data obtained from n = 3 zebrafish hearts per group. Error bars indicate mean ± s.d. Statistics were performed by two-tailed unpaired t-test.

Next, we assessed abundance and localization of Lamp1, a marker for late endosomes and lysosomes, in WT and *plna* R14del zebrafish hearts. *plna* R14del zebrafish hearts displayed a significantly increased number of Lamp1 foci compared to WT (**Figure 2C,D**), which is consistent with the observed increased expression of *lamp1b*. These Lamp1 foci frequently localized adjacent to Pln puncta near the nucleus (mean ± SD: 80.51% ± 14.60%) (**Figure 2C,E; Sup figure 3A,B**). This association of LAMP1 with PLN clusters was similarly observed in cardiomyocytes in LV tissue obtained from patients with end-stage cardiomyopathy carrying the PLN R14del variant (**Figure 2F**). In more detail, LAMP1 localized in a ring-like structure around the PLN clusters (**Figure 2F**). In contrast, the number of p62 foci, marking autophagosomes, was comparable between WT and *plna* R14del zebrafish hearts (**Sup figure 3C,D)**. Additionally, evaluation of autophagy flux did not show differences between WT and *plna* R14del zebrafish hearts (**Sup figure 3E,F**). Together, our results indicate that SR-derived vesicles in PLN R14del cardiomyocytes are surrounded by lysosomes suggesting that they are targeted for lysosomal degradation.

**Figure 3.**
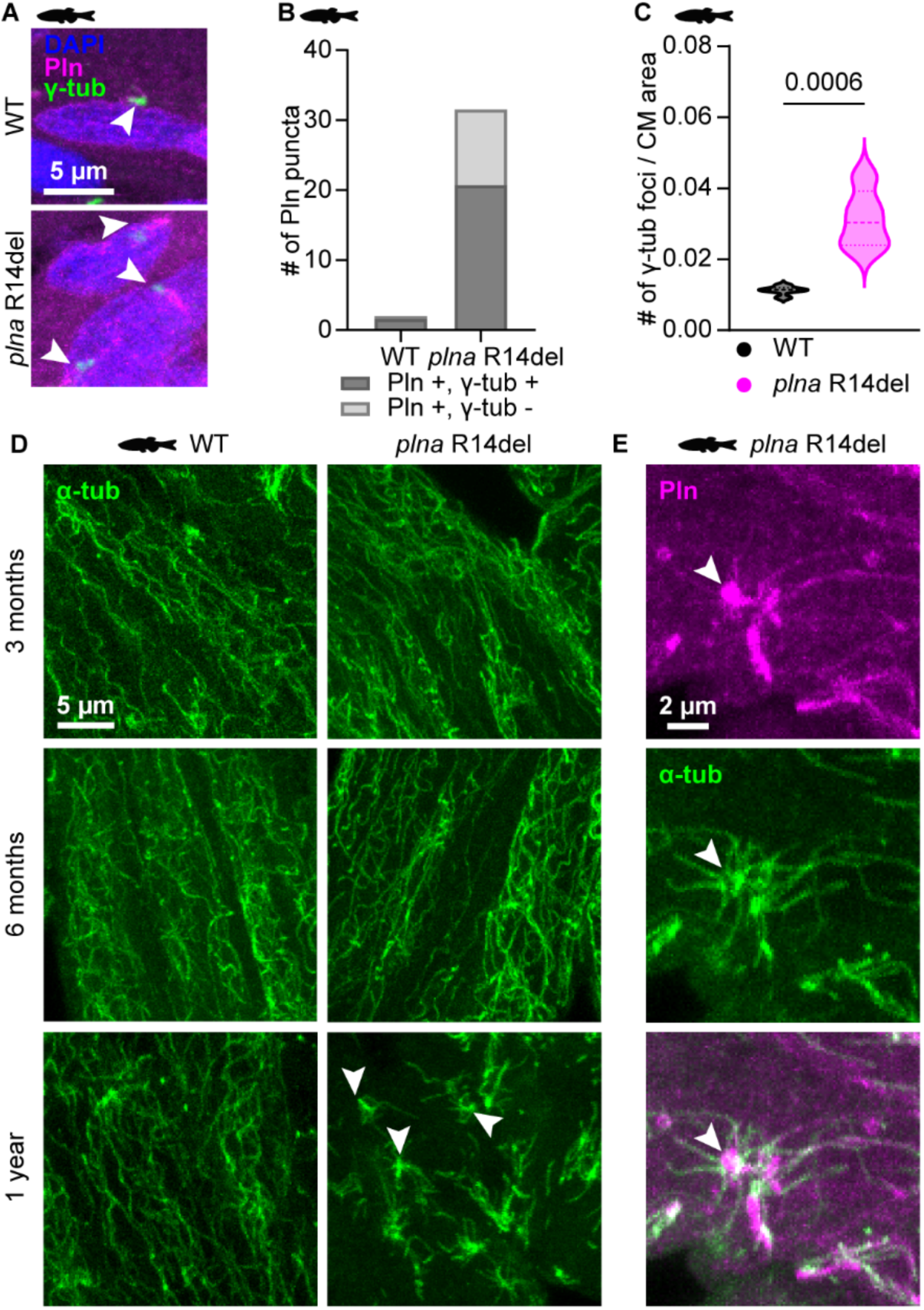
SR-derived vesicles localize at the MTOC inducing microtubule remodeling. **A.** Representative images of immunofluorescent staining against DAPI, Pln and γ-tubulin (γ-tub) in one year old wild type (WT) and *plna* R14del zebrafish. White arrowheads indicate γ-tubulin foci in WT hearts and γ-tubulin and Pln co-localization in *plna* R14del hearts. Data shown are representative for n = 8 WT and n = 8 *plna* R14del zebrafish hearts. Scale bar = 5 μm. **B.** Quantification of Pln puncta co-localized with γ-tubulin foci. Data obtained from n = 8 WT and n = 8 *plna* R14del zebrafish hearts. **C.** Quantification of number of γ-tubulin foci divided by cardiomyocyte (CM) area. Statistics were performed by two-tailed unpaired t-test. Data obtained from n = 8 WT and n = 8 *plna* R14del zebrafish hearts. **D.** Representative images of immunofluorescent staining against α-tubulin (α-tub) in three months, six months and one year old WT and *plna* R14del zebrafish hearts. Data shown are representative for at least n = 3 individuals per group per timepoint. Scale bar = 5 μm. White arrowheads indicate microtubule asters. **E**. Representative image of immunofluorescent staining against Pln and α-tub in one year old *plna* R14del zebrafish hearts. White arrowhead shows co-localization of Pln with a microtubule aster sprouting from the Pln puncta. Data shown are representative for n = 8 *plna* R14del zebrafish hearts. Scale bar = 2 μm.

### Ca^2+^ dysregulation impairs lysosomal function and promotes SR-derived vesicle accumulation

To further characterize the Lamp1 foci, we performed immunoelectron microscopy for Lamp1 in WT and *plna* R14del zebrafish hearts. In *plna* R14del hearts, more Lamp1^+^ organelles accumulated and these organelles predominantly corresponded to late endosomes/multivesicular bodies as well as enlarged endo-lysosomal compartments containing intraluminal vesicles and undegraded material, features that were not observed in WT hearts (**Figure 2G**). These ultrastructural characteristics indicate a slowed or hampered degradative rate and are consistent with a defect in endo-lysosomal progression, potentially reflecting impaired endosome-to-lysosome maturation in *plna* R14del hearts. Based on these observations and prior reports showing that Ca²⁺ overload disrupts lysosomal function in neurons (Jung et al., 2019; Kim et al., 2023; Mustaly-Kalimi et al., 2022), we hypothesized that elevated cytosolic Ca^2+^ in *plna* R14del hearts impairs the maturation of late endosomes into functional lysosomes. Supporting this hypothesis, we observed a reduction in DQ red BSA foci, a marker for functional lysosomes (Marwaha & Sharma, 2017), in cardiac slices from *plna* R14del zebrafish compared to WT (**Figure 2H,I**) as well as in thapsigargin treated cardiac slices (**Figure 2J,K**). To distinguish lysosomal dysfunction from reduced endocytic uptake, we performed co-uptake experiments using both DQ red BSA, marking functional lysosomes, and green BSA, reporting endocytosis rates. As the number of green BSA foci were unchanged between the samples, the data showed that the reduced DQ red BSA foci reflects impaired lysosomal function rather than reduced endocytic uptake (**Sup figure 3G-J**). These data clearly demonstrate an impaired endosome-to-lysosome maturation, lysosomal dysfunction, and incomplete degradation of endocytosed cargo in *plna* R14del mutants. This in turn induces a compensatory transcriptional program associated with endo-lysosomal biogenesis, similar to that observed in other molecular perturbations targeting the final stages of endo-lysosomal maturation (van der Beek et al., 2024); however, this response appears insufficient to restore effective lysosomal maturation and degradative capacity. Collectively, these results indicate that in PLN R14del cardiomyopathy, elevated cytosolic Ca²⁺ drives SR-derived vesicle formation, while persistent Ca²⁺ overload compromises lysosomal function together leading to progressive accumulation of these vesicles.

### SR-derived vesicles localize at the MTOC inducing microtubule remodeling

Currently, the underlying mechanism of the frequent perinuclear accumulation of SR-derived vesicles is unknown. Given their co-localization adjacent to lysosomes and the role of the microtubule organizing center (MTOC) as a key site for lysosomal clustering and clearance (Olzmann et al., 2008; Sasazawa et al., 2024), we hypothesized that SR-derived vesicles accumulate near the MTOC for lysosomal degradation. Consistent with this hypothesis, we observed frequent juxtaposed co-localization of Pln puncta with γ-tubulin foci, a marker for the MTOC (mean ± SD: 63.30% ± 15.30%) (**Figure 3A,B; Sup figure 4A,B**). Additionally, *plna* R14del hearts showed increased number of γ-tubulin foci (**Figure 3C**). Since accumulation of dynein-dynactin motor proteins at microtubule minus ends has been shown to induce microtubule aster formation (Tan et al., 2018), and given that SR-derived vesicles are likely associated with and retained at microtubule minus ends at the MTOC via motor protein interactions, we next examined the microtubule network in WT and *plna* R14del zebrafish hearts at different ages using α-tubulin as a microtubule marker. While α-tubulin organization in three and six months old *plna* R14del hearts showed no major differences compared to WT, hearts from one year old *plna* R14del zebrafish exhibited pronounced microtubule network remodeling characterized by microtubule aster formation (**Figure 3D**). High-resolution imaging revealed that these microtubule asters co-localize with Pln and sprout from Pln puncta (**Figure 3E**). Similarly, we identified characteristic microtubule aster structures forming around Pln puncta in thapsigargin treated WT cardiac slices (**Sup figure 4C**), suggesting that Ca^2+^ dysregulation alone is sufficient to induce SR-derived vesicle accumulation at the MTOC and drive microtubule remodeling. Notably, extensive microtubule network remodeling was also observed in LV tissue obtained from patients with end-stage cardiomyopathy carrying the PLN R14del variant, specifically in regions containing PLN clusters (**Sup figure 4D**). Together, these findings demonstrate that SR-derived vesicles accumulate at the MTOC, where they promote aberrant microtubule aster formation, leading to pathological remodeling of the microtubule network in PLN R14del cardiomyopathy.

**Figure 4.**
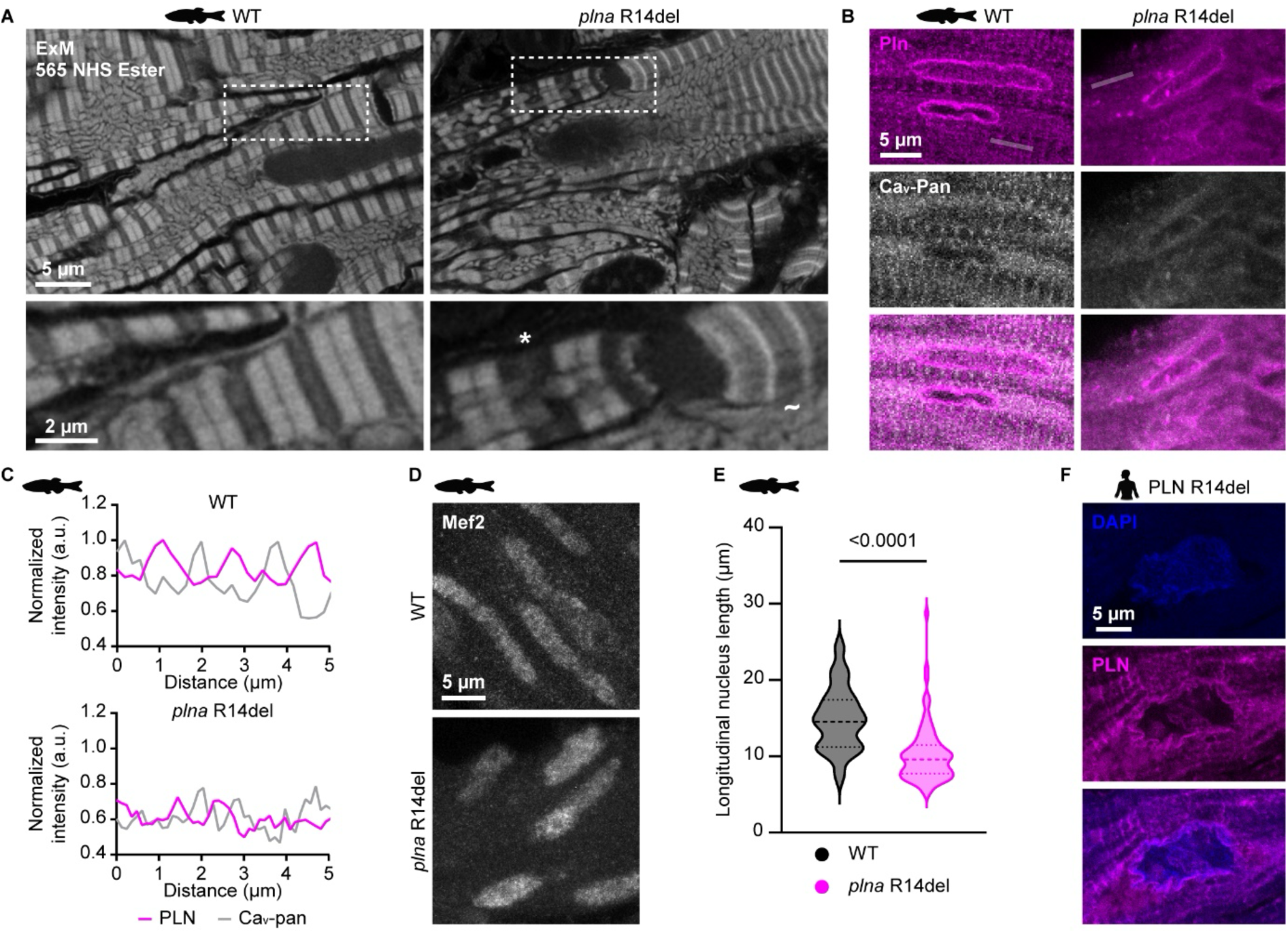
PLN R14del cardiomyocytes display structural disorganization. **A.** Representative images of 565 NHS ester general protein stain in one year old wild type (WT) and *plna* R14del expanded zebrafish hearts. * Indicates relaxed cardiomyocytes; ∼ indicates contracted cardiomyocytes. Data shown are representative for n = 3 WT and n = 3 *plna* R14del zebrafish hearts. Scale bar upper panel = 5 μm; scale bar lower panel = 2 μm; scale bars are corrected for pre-expansion dimensions. **B.** Representative images of immunofluorescent staining against Pln and Ca_v_-Pan in one year old WT and *plna* R14del zebrafish. Data shown are representative for n = 4 WT and n = 8 *plna* R14del zebrafish hearts. Scale bar = 5 μm. Line indicates measurement spot for panel C. **C.** Representative line plots showing Pln (magenta) and Ca_v_-Pan (grey) normalized intensity in WT and *plna* R14del cardiomyocytes. **D.** Representative images of immunofluorescent staining against Mef2 in one year old WT and *plna* R14del zebrafish. Data shown are representative for n = 6 WT and n = 7 *plna* R14del zebrafish hearts. Scale bar = 5 μm. **E.** Quantification cardiomyocyte longitudinal nucleus length in μm based on the Mef2 staining in one year old WT and *plna* R14del zebrafish hearts. Statistics were performed by two-tailed unpaired t-test. Data obtained from n = 6 WT and n = 7 *plna* R14del zebrafish hearts. **F.** Representative image of immunofluorescent staining against DAPI and PLN in left ventricular (LV) tissue obtained from patients with end-stage cardiomyopathy carrying the PLN R14del variant. Data shown are representative for n = 2 individuals. Scale bar = 5 μm.

### PLN R14del cardiomyocytes display structural disorganization

As microtubules play a central role in trafficking and spatial organization of mRNA, proteins, and organelles in cardiomyocytes (Uchida et al., 2022), we next assessed cellular and subcellular organization in one year old WT and *plna* R14del zebrafish hearts. To visualize cardiomyocyte architecture at high resolution, we established expansion microscopy (ExM) on thick zebrafish cardiac slices, combined with general protein staining using 565 NHS ester (Damstra et al., 2022). This approach enabled super-resolution imaging of sarcomere structures, including individual Z-lines, M-lines, A-bands, and I-bands (**Sup figure 5A,B**), as well as the spatial arrangement of mitochondria between sarcomeres (**Figure 4A**). In WT hearts, cardiomyocytes displayed highly organized sarcomeric alignment with uniformly spaced Z-lines (**Figure 4A; Sup video 1**). In contrast, *plna* R14del hearts exhibited irregular sarcomere organization and reduced alignment of contractile structures (**Figure 4A; Sup video 1**). Quantification of Z-line spacing revealed that relaxed sarcomeres in *plna* R14del cardiomyocytes had significantly increased inter-Z-line distances (**Sup figure 5C,D**), indicating a diastolic defect. Moreover, while WT cardiomyocytes showed uniform sarcomere states, either fully relaxed or contracted, *plna* R14del cardiomyocytes displayed both contracted and relaxed states within the same cell, suggesting uncoordinated contraction-relaxation (**Figure 4A; Sup video 1**). Given that arrhythmias are a hallmark of PLN R14del cardiomyopathy (van der Zwaag et al., 2013; van Rijsingen et al., 2014) and that microtubules facilitate ion channel trafficking to specific membrane domains (Uchida et al., 2022), we next assessed the localization of the L-type Ca^2+^ channels (LTCC) using Ca_v_-Pan as a pan-LTCC marker. In physiological conditions, LTCCs localize to cardiac dyads formed between the plasma membrane and the SR membranes, where they regulate Ca^2+^ influx during excitation-contraction coupling (Nguyen et al., 2023). In WT hearts, Pln and Ca_v_-Pan exhibited a highly ordered alternating striated pattern corresponding to the M- and Z-line sarcomeric architecture, respectively, which was lost in *plna* R14del hearts (**Figure 4B,C**). Additionally, as the microtubule network also protects the nucleus against mechanical stress during cardiomyocyte contraction (Ramdas & Shivashankar, 2015), we next examined nuclear shape. Consistent with microtubule disorganization observed in *plna* R14del hearts, we found altered nuclear morphology in cardiomyocytes. In more detail, cardiomyocyte nuclei, marked with Mef2, appeared more rounded in *plna* R14del zebrafish hearts compared to WT (**Figure 4D,E)**. Furthermore, LV tissue obtained from patients with end-stage cardiomyopathy carrying the PLN R14del variant exhibited severely deformed, irregularly shaped nuclei (**Figure 4F**). Taken together, these findings show that PLN R14del cardiomyocytes display widespread structural disorganization, including sarcomere misalignment, mislocalization of LTCCs, and altered nuclear morphology. These abnormalities are likely driven by concurrent microtubule network disruption and may underly the impaired contractility and arrhythmogenic phenotype characteristic of PLN R14del cardiomyopathy.

**Figure 5.**
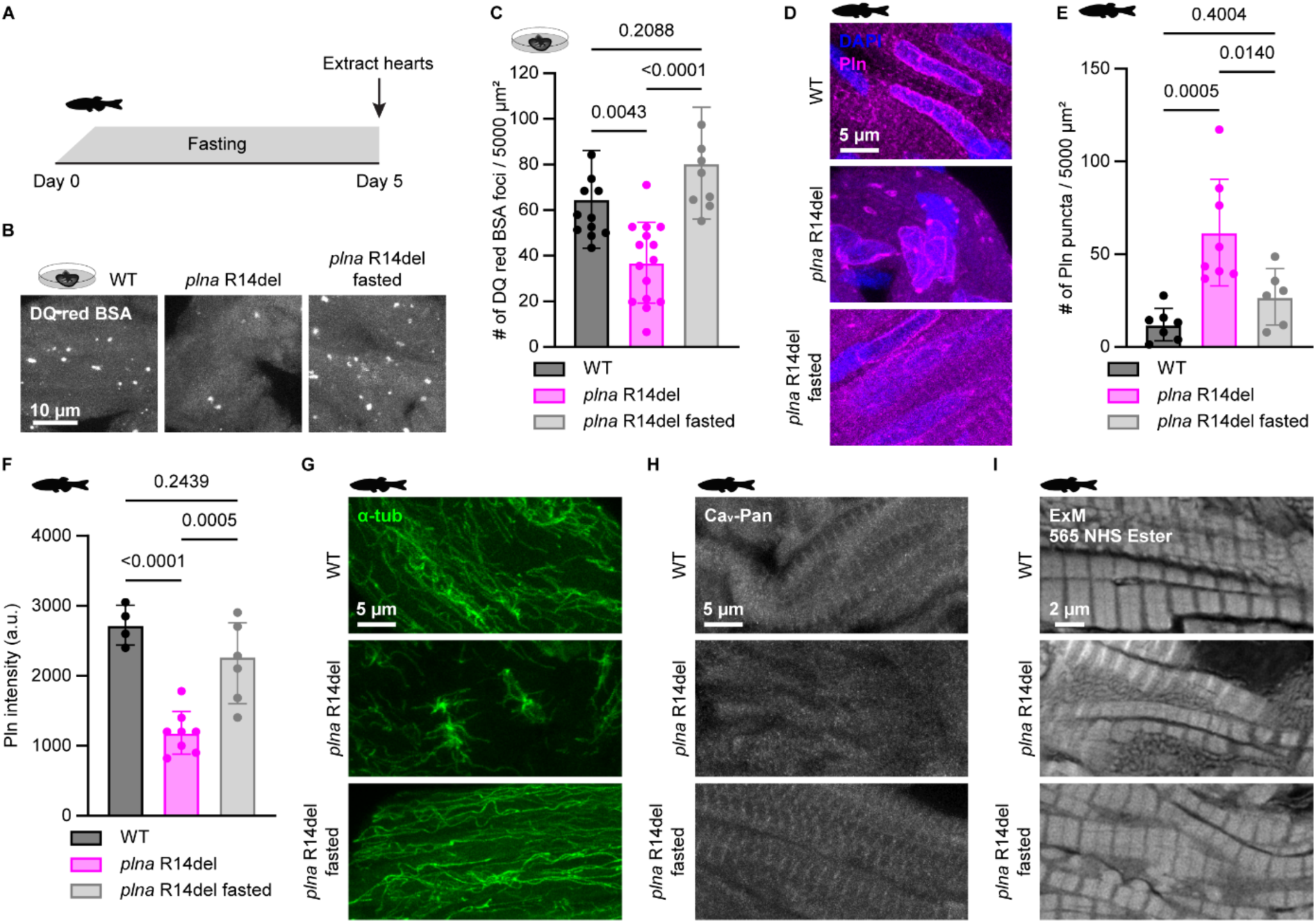
Fasting restores lysosomal function and rescues SR-derived vesicle accumulation, microtubule network integrity and cellular organization in *plna* R14del zebrafish hearts. **A.** Workflow fasting in *plna* R14del one year old zebrafish. **B.** Representative images of DQ red BSA *ex vivo* assay in one year old wild type (WT) and fasted versus non-fasted *plna* R14del hearts. Dots represent functional lysosomes. Data shown are representative for at least n = 3 per group. Scale bar = 10 μm. **C.** Quantification of number of DQ red BSA foci per 5000 μm^2^. Data points represent an individual measurement per image; multiple points originate from one individual; at least n = 3 individuals were included in each group. Error bars indicate mean ± s.d. Statistics were performed by one-way ANOVA. **D.** Representative images of immunofluorescent staining against DAPI and Pln in one year old WT and non-fasted versus fasted *plna* R14del zebrafish. Data shown are representative for at least n = 6 per group. Scale bar = 5 μm. **E-F** Quantification of Pln puncta per 5000 μm^2^ **(E)** and overall Pln intensity **(F)** in one year old WT and non-fasted versus fasted *plna* R14del zebrafish. Data points represent averages of individual hearts with at least n = 4 per group. Error bars indicate mean ± s.d. Statistics were performed by one-way ANOVA. **G.** Representative images of immunofluorescent staining against α-tubulin (α-tub) in one year old WT and non-fasted versus fasted *plna* R14del zebrafish. Data shown are representative for at least n = 3 per group. Scale bar = 5 μm. **H.** Representative images of immunofluorescent staining against Ca_v_-Pan in one year old WT and non-fasted versus fasted *plna* R14del zebrafish. Data shown are representative for at least n = 3 per group. Scale bar = 5 μm. **I.** Representative images of 565 NHS ester general protein stain in one year old WT and non-fasted versus fasted *plna* R14del expanded zebrafish hearts. Data shown are representative for at least n = 3 per group. Scale bar = 2 μm; scale bar is corrected for pre-expansion dimensions.

### Fasting restores lysosomal function and reduces SR-derived vesicle accumulation in *plna* R14del zebrafish hearts independent of macro-autophagy

Thus far, our results demonstrate that persistent Ca^2+^ dysregulation and subsequent lysosomal dysfunction drive PLN R14del disease progression through perinuclear PLN accumulation and cellular remodeling. Given that nutrient deprivation is a well-established activator of lysosomal function and has been shown to reduce protein aggregation in degenerating neurons (Ma et al., 2019; McGuire & Forgac, 2018; L. Wu et al., 2025; Young et al., 2009), we next investigated whether fasting could rescue or attenuate the accumulation of SR-derived vesicles in *plna* R14del zebrafish by reactivating lysosomal clearance. To this end, we subjected *plna* R14del zebrafish to a five-day fasting regime aimed at reactivating lysosomal pathways (**Figure 5A**). Excitingly, fasting effectively restored lysosome function in *plna* R14del hearts, as evidenced by the number of DQ red BSA foci in *plna* R14del restoring to WT levels (**Figure 5B,C**). Strikingly, fasting robustly reduced the number of Pln puncta in *plna* R14del hearts, restoring them to WT levels (**Figure 5D,E**). Furthermore, fasting improved Pln localization patterns in *plna* R14del hearts to a distribution pattern resembling physiological SR and NE organization as seen in WT cardiomyocytes (**Figure 5D-F**). To determine whether fasting reduces SR-derived vesicles accumulation solely through lysosomal reactivation or also involves upregulation of macro-autophagy in our model system, *plna* R14del zebrafish were treated with rapamycin, an mTOR inhibitor known to induce macro-autophagy (Chávez et al., 2020). *plna* R14del zebrafish treated with rapamycin showed no reduction in Pln puncta, suggesting that fasting rescues SR-derived vesicle accumulation through a macro-autophagy independent mechanism (**Sup figure 6A-C**). In line with this, fasting did not stimulate autophagy flux in WT nor *plna* R14del zebrafish hearts (**Sup figure 6D-F**). Additionally, p62 foci, marking autophagosomes, did not show differences in localization before and after fasting in WT nor *plna* R14del zebrafish (**Sup figure 6G**). Together, these results demonstrate that fasting reverses SR-derived vesicle accumulation and restores physiological Pln localization in *plna* R14del zebrafish hearts through direct stimulation of lysosomal function independent of macro-autophagy.

### Fasting restores the microtubule network integrity and cellular organization in *plna* R14del zebrafish hearts

Given that SR-derived vesicle accumulation in *plna* R14del hearts is associated with microtubule network disruption and cellular disorganization and that fasting reduces the accumulation of aberrant SR structures in *plna* R14del hearts, we next assessed whether fasting could also restore the microtubule network and overall cellular organization. Excitingly, fasting markedly restored microtubule network organization in one year old *plna* R14del zebrafish: instead of short aster-like structures clustered around the SR-derived vesicles, we observed a homogeneous and dense microtubule network resembling the organization seen in WT hearts (**Figure 5G**). To determine whether microtubule restoration also re-established correct localization of ion channels, we examined the distribution of LTCCs marked by Ca_v_-Pan. We observed that LTCCs regained their characteristic striated pattern in fasted *plna* R14del hearts (**Figure 5H**). Finally, we assessed sarcomere organization in fasted *plna* R14del hearts using ExM and observed that fasting restored sarcomere alignment to a pattern comparable to WT hearts, indicating improved contractile organization (**Figure 5I**). Collectively, these results show that lysosomal reactivation through fasting can reverse key disease-causing mechanisms in *plna* R14del cardiomyopathy progression, including Pln^+^ SR-derived vesicle accumulation, microtubule network integrity, ion channel localization, and overall cellular architecture, at a clinically relevant disease stage. This reveals a direct mechanistic link between lysosomal function, cytoskeletal organization, and cardiomyocyte structural integrity, which opens new avenues for therapeutic interventions in PLN R14del cardiomyopathy.

## Discussion

In this study, we uncover the mechanistic origin and pathological consequences of the previously described PLN aggregates in PLN R14del cardiomyopathy. Using a combination of *plna* R14del zebrafish, LV tissue obtained from patients with end-stage cardiomyopathy carrying the PLN R14del variant, and pharmacological modeling in WT cardiac slices, we propose a three-step model linking Ca^2+^ dysregulation to structural remodeling: (1) elevated cytosolic Ca^2+^, due to impaired SERCA2a activity, promotes aberrant formation of PLN-containing SR-derived vesicles that localize adjacent to lysosomes; (2) persistent Ca^2+^ overload impairs lysosomal function, leading to progressive accumulation of PLN-containing vesicle; and (3) vesicle accumulation at the MTOC disrupts the microtubule network, which in turn triggers nuclear deformation, sarcomere misalignment, and broader cellular disorganization. Importantly, key disease phenotypes, including SR-derived vesicle accumulation, lysosomal dysfunction, and microtubule remodeling, were consistently observed across *plna* R14del zebrafish, LV tissue obtained from patients with end-stage cardiomyopathy carrying the PLN R14del variant, and thapsigargin-treated WT cardiac slices, strengthening the relevance of the model in a disease progression context. Finally, we demonstrate that lysosomal reactivation through fasting reverses these pathological features at a clinically relevant stage in our humanized *plna* R14del zebrafish model, highlighting SR-derived vesicle accumulation at the MTOC as a central driver of the disease and pointing to lysosomal reactivation as a promising therapeutic strategy to halt or even reverse PLN R14del cardiomyopathy progression.

Abnormal PLN protein accumulation is a consistent pathological hallmark across all PLN R14del model systems, yet the mechanisms that drive this accumulation and its pathological relevance remain unresolved (Stege et al., 2024). Our data demonstrate that Ca²⁺ overload in *plna* R14del cardiomyocytes disrupts lysosomal function and drives the accumulation of SR-derived vesicles, supporting the model in which the R14del variant acts as a super-inhibitor for SERCA2a and thereby perturbs intracellular Ca²⁺ homeostasis (Haghighi et al., 2006, 2021; Kamel et al., 2021; Kumar et al., 2023; Vafiadaki et al., 2022). This interpretation aligns with a recent study showing that PLN R14del-mediated Ca²⁺ dysregulation impairs autophagosome-lysosome fusion in HEK293 and H9c2 cells, resulting in PLN protein accumulation, further strengthening the link between Ca²⁺ imbalance and defective lysosomal processing (Vafiadaki et al., 2024). In contrast, other reports propose that PLN R14del partially loses its inhibitory control over SERCA2a, causing enhanced SR Ca²⁺ uptake (Cleary et al., 2025; Maniezzi et al., 2024; Stege et al., 2023). Importantly, impaired lysosomal clearance has been observed under both Ca^2+^ overload and Ca^2+^ depletion (Fedeli et al., 2019; Jung et al., 2019; Kim et al., 2023; Mustaly-Kalimi et al., 2022), suggesting that deviations in either direction may converge on a shared pathological endpoint: accumulation of aberrant SR-derived vesicles due to lysosomal dysfunction. Together, these findings support the view that lysosomal dysfunction, initiated by disturbed Ca²⁺ homeostasis rather than a single directional change, is a central mechanism driving PLN accumulation and contributing to disease progression in PLN R14del cardiomyopathy. Notably, lysosomal dysfunction has been most clearly associated with the development of various forms of cardiomyopathies in patients carrying mutations in lysosomal genes such as *LAMP2*, *CTSL*, and *GAA* (Felice, 2017; Stypmann et al., 2002; Yang et al., 2005), highlighting the critical role of defective lysosomal processing in cardiomyopathy onset and progression.

Interestingly, although we identify lysosomal dysfunction as a central driver of PLN R14del cardiomyopathy onset and progression, the accumulation of PLN-containing SR-derived vesicles does not appear to involve classical macro-autophagy, indicating that these structures bypass canonical autophagosome-mediated degradation. Instead, their clearance may depend on alternative quality-control mechanisms, potentially including ER-to-lysosome-associated degradation (ERLAD), a pathway recently identified for the selective removal of aberrant ER membrane proteins (Hayashi et al., 2023; Salomo-Coll et al., 2025). Collectively, these observations suggest that PLN vesicle accumulation reflects disruption of a specialized lysosome-dependent process rather than a general defect in macro-autophagy.

In *plna* R14del zebrafish, Ca^2+^ cycling abnormalities emerge as early as three days post fertilization (Kamel et al., 2021), yet the accumulation of SR-derived vesicles becomes apparent only from approximately one year of age. This striking temporal gap suggests that vesicle accumulation is not solely a consequence of Ca^2+^ mediated lysosomal dysfunction, but instead reflects the interplay between early Ca^2+^ dysregulation and an additional factor. Such a factor could be the decline in lysosomal function upon ageing (Colacurcio & Nixon, 2016; Lim et al., 2024; Nixon, 2020; Triolo & Hood, 2021). As ageing is accompanied by a progressive reduction in lysosomal efficiency, we propose a model in which early Ca²⁺ dysregulation-induced lysosomal impairment places the system near, but not beyond, a functional threshold. Over time, physiological lysosomal decline may tip this balance, rendering clearance insufficient and allowing SR-derived vesicles to accumulate. This framework aligns with the concept that mild lysosomal defects remain subclinical until ageing-associated deterioration further compromises lysosome function (Colacurcio & Nixon, 2016). Supporting the broader link between lysosomal function and ageing, recent work shows that modulating V-ATPase activity, a proton pump essential for lysosomal acidification, is tightly associated with cellular ageing (Arif et al., 2025). Moreover, enhanced V-ATPase activity expands lifespan in *C. elegans* (Baxi et al., 2017), whereas mutations in V-ATPase subunits are associated with ageing and neurodegenerative disease (Colacurcio & Nixon, 2016). Together, these observations support a model in which Ca^2+^ dysregulation initiates the formation of SR-derived vesicles in early disease stages, but efficient lysosomal clearance in young cardiomyocytes prevents their accumulation. As lysosomal capacity naturally declines with age, this clearance becomes progressively compromised, enabling SR-derived vesicles to build up and drive downstream pathological remodeling.

Importantly, our data indicates that microtubule remodeling is a critical downstream consequence of Ca²⁺ dysregulation and lysosomal impairment in PLN R14del cardiomyocytes. The cardiac microtubule network is a central organizer of mechanics, Ca²⁺ handling, and protein trafficking: it regulates viscoelasticity, dyadic organization, and delivery of ion channels and junctional proteins to the membrane (Caporizzo & Prosser, 2022; Uchida et al., 2022). In heart failure and inherited cardiomyopathies, microtubule densification and altered post-translational modification have been linked to increased cellular stiffness, impaired contractility, and defective trafficking of key structural and signaling complexes (Algül et al., 2023; Caporizzo & Prosser, 2022). Consistently, genetic disruption of microtubule-associated proteins such as CENP-F leads to DCM, underscoring that primary defects in microtubule organization are sufficient to drive DCM-like remodeling (Dees et al., 2012). At the intercalated disc, microtubules are required for proper delivery of connexin-43 and interact functionally with desmosomal complexes. Perturbation of these interactions promotes desmosome and gap-junction remodeling and creates an arrhythmogenic substrate characteristic of ACM (Chkourko et al., 2012; Costa et al., 2021; Delmar & McKenna, 2010; Macquart et al., 2019; Patel et al., 2014). In this context, microtubule remodeling provides a unifying mechanistic framework to explain why the same PLN R14del variant can lead to divergent cardiomyopathy phenotypes across patients (Hof et al., 2019; Stege et al., 2024; Vafiadaki et al., 2023; van der Zwaag et al., 2012). Rather than inducing uniform downstream effects, microtubule remodeling triggered by Ca²⁺ dysregulation and lysosomal dysfunction may stochastically bias cardiomyocytes toward distinct structural states. In some cardiomyocytes, microtubule defects may predominantly compromise intercalated-disc integrity and junctional protein localization - consistent with reduced plakoglobin at cell-cell junctions observed in PLN R14del patients with ACM features (Te Rijdt et al., 2019) - whereas in others, microtubule alterations may primarily affect sarcomeric organization, cellular stiffness, and excitation-contraction coupling, resulting in a DCM phenotype with preserved junctional architecture. This variability may provide a mechanistic explanation for the marked phenotypic heterogeneity observed across patients carrying the PLN R14del variant, as well as for the coexistence of overlapping ACM and DCM features within a single patient (te Rijdt et al., 2016).

Strikingly, we show that lysosomal reactivation by fasting rescues the pathological remodeling in *plna* R14del zebrafish hearts. Moreover, fasting reduced SR-derived vesicle accumulation, re-established a homogenous and organized microtubule network, restored striated localization of LTCCs at dyad structures and normalized sarcomere alignment in *plna* R14del zebrafish hearts. These findings position lysosomal reactivation as a central mechanism capable of halting or even reversing PLN R14del disease progression at a clinically relevant stage. Consistent with this concept, repetitive stimulation of the autophagy-lysosome machinery by intermittent fasting in mice preconditions the myocardium against ischemia-reperfusion injury and improved cardiac function in patients with myocardial infarction (Dutzmann et al., 2024; Godar et al., 2015), underscoring the translational relevance of this approach in human patients. As a dietary intervention, fasting may allow more rapid clinical implementation than pharmacological strategies that require extensive development and regulatory approval. Apart from lysosomal activation through fasting, direct activation of lysosomal biogenesis also promotes cardioprotection: forced activation of the lysosomal transcription factor TFEB in cardiac proteinopathy models enhances autophagy-lysosome function and ameliorates proteotoxic cardiomyopathy, while pharmacological TFEB activation with agents such as trehalose improves post-infarction remodeling and reduces fibrosis (Ma et al., 2019; Pan et al., 2017; Sciarretta et al., 2018). It is important to note that while lysosomal activation represents a promising therapeutic avenue, its implementation must be carefully controlled. Translation to clinical settings should proceed cautiously, as excessive or prolonged enhancement of the lysosome system could compromise cellular homeostasis and potentially lead to adverse effects. Nevertheless, together with our zebrafish data, these studies support a model in which boosting lysosomal function, whether by dietary interventions or pharmacological lysosome activators, can reverse structural and functional abnormalities and may represent a promising therapeutic avenue for PLN R14del cardiomyopathy.

## Acknowledgements

We thank the Hubrecht Animal facility, the Hubrecht imaging center and the Hubrecht FACS facility for their technical assistance; J. Korving and H. Begthel for their histology assistance; E.A. Katrukha, R.R. Padala and A. Griffa for bioinformatic analysis and data visualization; T.P. de Boer and Y. Onderwater for technical assistance. We acknowledge the UNRAVEL team for patient inclusion and tissue collection. We also thank A. Akhmanova, A.K. Serweta and A.R. Keijzer for their valuable advice and recommendations. Furthermore, we thank E. Penning de Vries, M.J.C. Houtman, I.R. Kelters, C.N. van der Wilt and C. Snijders Blok for their assistance in collecting human patient material. Additionally, we thank F. Tessadori, M. Bouwman, L. Florit Gonzalez and D.E.M. de Bakker for their careful proofreading and valuable suggestions. Lastly, we thank all members of the Bakkers group for their insightful feedback and engaging discussions. This work was supported by ZonMw open competition (I.G., M.H., & J.B.), ZonMW Clinical Fellow (A.S.J.M.R.) and ERC HORIZON IMPACT (A.S.J.M.R.). Additionally, part of the work has been enabled by the NEMI and NL-BioImaging infrastructures sponsored by NWO National Roadmap for Large-Scale Research Infrastructure (NWO 184.034.014 and NWO 184.036.012).

## Methods

### Animal experiments

All zebrafish experiments were conducted under the guidelines of the animal welfare committee of the Royal Netherlands Academy of Arts and Sciences (KNAW) and were approved by the local ethics committee at the KNAW. All adult zebrafish lines were kept at stock density of max 7 fish per liter with the following parameters: temperature around 28 °C, 14 hours light / 10 hours dark cycle, pH around 7.5 and conductivity around 450 uS/cm. Adult zebrafish (> 14 weeks) were fed two times a day with flaks containing TetraMin and living artemia, with the exception of zebrafish part of the fasting experiments in which fish were deprived from food for five days.

### Generation of zebrafish lines

The following zebrafish lines were used: TL, *plna* R14del/R14del;*plnb* +/+ zebrafish (referred to as *plna* R14del zebrafish) (Tessadori et al., 2018), Tg(*myl7*:GCaMP6f-nls-T2A-RCaMP107-NES-pA)^hu11799^ (Bertozzi et al., 2021), Tg(*myl7:actn3b*-EGFP) (Lin et al., 2012), Tg(*myl7*:mKate-KDEL;exorh:GFP) and Tg(*myl7*:PercevalHR). Both males and females were used for all zebrafish experiments. The Tg(*myl7*:mKate-KDEL;*exorh*:GFP) line was generated by combining p5E *myl7* (Nguyen et al., 2023), pME mKate-KDEL (KDEL sequence = atgctgctatccgtgccgttgctgctcggcctcctcggcctggccgtcgccctcgag), p3E polyA (Kwan et al., 2007) and pDest *exorh*:GFP (gift from Christian Mosimann (Kemmler et al., 2023) (Addgene plasmid # 195983; http://n2t.net/addgene:195983; RRID:Addgene_195983)) and injecting the construct with Tol2 transpoase. The Tg(*myl7*:PercevalHR) line was generated by combining p5E *myl7* (Nguyen et al., 2023), pME PercevalHR (GW1-PercevalHR was a gift from Gary Yellen (Tantama et al., 2013) (Addgene plasmid # 49082; http://n2t.net/addgene:49082; RRID:Addgene_49082)), p3E polyA (Kwan et al., 2007) and pDestpA4 (Kwan et al., 2007) and injecting the construct with Tol2 transpose.

### Heart collection for histological analysis

In zebrafish, adult zebrafish ventricles were isolated in extraction buffer (PBS, 10% KCL (BOOM, 7447-40-7), heparin (Sigma-Aldrich, 9041-08-1)) and fixed in 4% PFA + 0.1% glutaraldehyde (Sigma-Aldrich, 111-30-8) in PBS (37 °C, 30 min). Subsequently, the hearts were washed 2× 10 min in NaBH_4_ (0.1 M) (Sigma-Aldrich, 16940-66-2) in PBS followed by 1x 30 min glycine (ThermoFisher Scientific, 56-40-6) in PBS (0.1 M). Next, the hearts were washed 3× 10 min in 4% sucrose (Sigma-Aldrich, 57-50-1) in PBS followed by 30% sucrose (Sigma-Aldrich, 57-50-1) O/N on a roller bank at 4 °C. Next, the hearts were embedded in tissue freezing medium (Leica, 03829006). Cryo-sectioning of the hearts was performed at 10 µm thickness.

For human tissue, LV tissue was obtained from six PLN R14del patients who underwent heart transplantation at the University Medical Center Utrecht. Patients provided written informed consent. Collection and use of patient material was approved by the institutional review board (UCC-UNRAVEL, TCBio #12-387) and compliant with the declaration of Helsinki. For original evaluation of PLN localization, fixed FFPE biopsies (n=2) were used. For further examination of lysosomes and microtubules, fresh LV tissue was used (n=4). In more detail, the fresh cardiac tissue was fixed in 4% PFA + 0.1% glutaraldehyde (Sigma-Aldrich, 111-30-8) in PBS (37 °C, O/N). The next morning, the cardiac tissue was washed 2× 10 min in NaBH_4_ (0.1 M) (Sigma-Aldrich, 16940-66-2) in PBS followed by 1x 30 min glycine (0.1 M) (ThermoFisher Scientific, 56-40-6) in PBS. Subsequently, the cardiac tissues were dehydrated in ethanol series and embedded in paraffin. The paraffin blocks were sectioned at 10 µm thickness.

### Immunohistochemistry

On zebrafish cryosections, antigen retrieval was performed by heating slides containing heart sections at 85 °C in 10 mM sodium citrate (Merck, 1.06431) buffer (pH 6) for 15 min. For human paraffin sections, antigen retrieval was performed by heating slides containing heart sections under pressure at 120 °C in 10 mM sodium citrate (Merck, 1.06431) buffer (pH 6) for 1 hour. The following primary antibodies were used across all samples: PLN (1:100, Invitrogen, MA3-922), γ-TUB (1:400, Sigma-Aldrich, T6557), LAMP1 (1:50, Abcam, ab24170), α-TUB (1:200, Sigma-Aldrich, T5168), Ca_v_-Pan (1:200, Alomone labs, ACC-004), Mef2 (1:500, Boster Biological Technology, DZ01398-1), GRP78 (1:200, Proteintech, 11587-1-AP) and p62 (SQSTM1) (1:500, MBL International, PM045MS). Secondary antibodies included anti-mouse IgG1 (1:500, Alexa 488, ThermoFisher Scientific), anti-mouse IgG2a (1:500, Alexa 568, ThermoFisher Scientific; 1:500, Alexa 633, ThermoFisher Scientific) and anti-rabbit (1:500, Cy2, Jackson Laboratories; 1:500, Cy5, Jackson Laboratories). DAPI (1:1000, ThermoFisher Scientific) was used as nuclear marker. Images of immunofluorescence were acquired using a Sp8 microscope (Leica) or a STELLARIS (Leica). Picrosirius red staining was performed as described before (Kamel et al., 2021) and sections were imaged using a VS200 Slide Scanner (Olympus).

### Immunohistochemistry quantifications

Immunohistochemistry quantifications were all performed using Fiji. For Pln puncta quantification, a max projection of the z-stacks was made and three different regions of 3817.66 µm² from each individual were included per genotype or treatment. Subsequently, averages per individual were plotted. For Pln intensity, a sum of the z-stacks was make and a 10 pixel width line was drawn perpendicular to the striation stretching inside the nucleus. The highest peak was used as NE intensity, while the following peak was used for the SR intensity. Multiple intensities were measured per individual and plotted. Number of γ-tubulin, Lamp1 and p62 foci were quantified in max projections of the z-stacks from two different regions of 3817.66 µm² from each individual using the threshold and analyze particles function in Fiji. The number of particles was then divided by total cardiomyocyte area determined by another threshold layer. For Lamp1 co-localization with Pln, max projections of the z-stacks from two different regions of 3817.66 µm² from each individual were included, in which both the Lamp1 foci and Pln puncta were manually highlighted. Subsequently, when Lamp1 foci and Pln puncta were adjacent to each other they were counted as co-localized. Lastly, the number of Pln puncta co-localized with Lamp1 foci was plotted. This same quantification was done to determine γ-tubulin foci and Pln puncta co-localization. To quantify nucleus length, max projections of z-stacks from two different regions of 3817.66 µm² from each individual were used. Only Mef2 staining in cardiomyocytes which were cut longitudinal were included and the longest axis was measured. Multiple nuclei were quantified and plotted. For line plots, a 10 pixel width plot line was drawn perpendicular to the striation pattern (Pln & Ca_V_-Pan) and along the co-localization axis (Pln & Lamp1 + Pln & γ-tubulin). Intensities were normalized by dividing the values by the highest value within that same plot line.

### *Ex vivo* cardiac slices treatments

Adult zebrafish ventricles were isolated in extraction buffer (culture medium, 10% KCL (BOOM, 7447-40-7), heparin (Sigma-Aldrich, 9041-08-1)). Culture medium consisted of 120 mM NaCl (Sigma-Aldrich, 7647-14-5), 3 mM KCl (BOOM, 7447-40-7), 2 mM CaCl_2_ (J.T. Baker, 10035-04-8), 1 mM MgCl_2_ (BOOM, 76021126), 3 mM NaHCO_3_ (Sigma-Aldrich, 144-55-8), 1.25 mM NaH_2_PO_4_ (J.T. Baker, 1347-23-5), 15 mM HEPES (Sigma-Aldrich, 7365-45-9), 5 mM glucose (Sigma-Aldrich, 50-99-7), 0.2 mM pyruvate (Sigma-Aldrich, 127-17-3), 0.5 mM GlutaMax (Gibco, 35050-038) (pH 7.4). Hearts were mounted in 1% agarose (Invitrogen, 16500-500) in 7×7×5 basemolds (VWR, 720-0820) and slices were made using a vibratome (150 μm) (Leica VT 1200S). For Ca^2+^ cycling experiments, Tg(*myl7*:GCaMP6f-nls-T2A-RCaMP107-NES-pA)^hu11799^ fish were used and slices were mounted in 0.5% agarose (Invitrogen, 16500-500). Subsequently, the slices were treated for 5 hours at 28 °C with either culture medium alone or thapsigargin (100 μM) (Biotechne, 67526-95-8). After 5 hours, the hearts were paced at 0.5 Hz and 28 °C (Nguyen et al., 2023). Videos were obtained by using the THUNDER microscope (Leica). Data analysis was performed as previously described (Nguyen et al., 2023). For rescue experiment with thapsigargin and BAPTA-AM, heart slices (150 μm) were treated for 5 hours at 28 °C with either culture medium, thapsigargin (100 μM) (Biotechne, 67526-95-8) or thapsigargin (100 μM) (Biotechne, 67526-95-8) in combination with BAPTA-AM (100 μM) (MedChemExpress, 126150-97-8). After treatment slices were fixed in 4% PFA in PBS (37 °C, 30 min). Next, heart slices were washed 3× 10 min in 4% sucrose (Sigma-Aldrich, 57-50-1) in PBS followed by 30% sucrose (Sigma-Aldrich, 57-50-1) O/N on a roller bank at 4 °C. Next, the hearts were embedded in tissue freezing medium (Leica, 03829006) and 10 μm thick cryosections were made which were used for immunohistochemistry. For DQ red BSA, 1 mg DQ red BSA (MedChemExpress, HY-D2449) was dissolved in 1 ml PBS. Regarding green BSA, 1% stock solution was made in dH₂O. Next, cardiac slices (150 μm) were incubated with DQ red BSA (MedChemExpress, HY-D2449) 20 μg/ml or DQ red BSA (MedChemExpress, HY-D2449) 20 μg/ml together with green BSA (Invitrogen, A13100) 20 μg/ml for 1 hour at 28 °C in culture medium. After treatment, slices were fixed in 4% PFA in PBS (37 °C, 30 min). Subsequently, cardiac slices were washed 3× 10 min in 4% sucrose (Bio-connect, HY-D2449) in PBS followed by 30% sucrose (Bio-connect, HY-D2449) O/N on a roller bank at 4 °C. Next, the hearts were embedded in tissue freezing medium (Leica, 03829006) at 10 μm thick cryosections were made while keeping the material in the dark. The sections were imaged using a Sp8 microscope (Leica). To quantify DQ red foci, max projection of z-stacks from three different regions of 3817.66 µm² from each individual were included and foci were counted manually.

### Single cell sequencing

Ventricles from Tg(*myl7*:PercevalHR) fish were extracted and collected in HBSS Ca^2+^/Mg^2+^ free medium (Life Technologies, 14175053). Subsequently, five ventricles were pooled together and digested in HBSS Ca^2+^/Mg^2+^ free media (Life Technologies, 14175053) containing 0.1% collagenase type II (Gibco, 17101015) at 32°C for 30 min followed by 1x TrypLE Express (Gibco, 12605036) for 15-30 min at 32°C with agitation. Dissociated cells were then FACSorted on mVenus and subjected to single-cell mRNA-seq. SORT-seq was performed as described before (Honkoop et al., 2019), with the exception that reads were mapped against a genome reference using STAR. For further analysis we used poisson corrected count tables and we combined three 384-cell plates for both genotypes (WT and *plna* R14del). Based on ncount distribution in all the plates, we used a cutoff at minimally 1000 ncounts per cell before further analysis. This resulted in 449 WT and 576 *plna* R14del single cells. Batch-effects were analyzed and showed no plate-specific clustering. Differential gene expression analysis was performed using the Idents algorithm. Fold enrichment of downregulated and upregulated GO terms were identified using the DAVID online GO analysis tool. Genes with P values < 0.05 and a log₂(fold change) higher than +/− 0.7 were included.

### Expansion microscopy

#### Tissue collection

Adult zebrafish ventricles were isolated in extraction buffer (PBS, 10% KCL (BOOM, 7447-40-7), heparin (Sigma-Aldrich, 9041-08-1)) followed by fixation in 4% PFA + 0.1% glutaraldehyde in PBS (37 °C, 30 min). Subsequently, the hearts were washed 2× 10 min in NaBH_4_ (0.1 M) (Sigma-Aldrich, 16940-66-2) in PBS followed by 1x 30 min glycine (ThermoFisher Scientific, 56-40-6) in PBS (0.1 M). Next, hearts were mounted in 1% agarose (Invitrogen, 16500-500) in 7×7×5 basemolds (VWR, 720-0820) and slices were made using a vibratome (150 μm) (Leica VT 1200S). Cardiac slices were stored in PBS + 0.05% sodium azide at 4 °C until further use.

#### Heart section expansion

Fixed heart sections were processed using an ultrastructure expansion microscopy (U-ExM) according to a protocol adapted from (Gambarotto et al., 2021) with modifications to accommodate our samples. Briefly, samples were anchored in anchoring solution (PBS (Gibco, 18912-014), 1.4% formaldehyde (ThermoFisher Scientific, 28908), 3% acrylamide (Sigma-Aldrich, A4058)) O/N at 37°C, then washed in 3× 10min PBS. The anchored samples were incubated first in monomer solution (PBS, 19% sodium acrylate, 10% acrylamide, 0.1% bis-acrylamide (Bio-Rad, 1610142) and next in gelation solution (the monomer solution supplemented with 0.001% 4-Hydroxy-TEMPO (Sigma-Aldrich, 176141), 0.2% APS (Sigma-Aldrich, A3678 and 0.2% TEMED (BioRad, 1610800)). Both of the incubation steps were performed on ice, with shaking; the former for 1 hour, the latter for 30 min. Subsequently, the sections were mounted in gelation chambers, there embedded in the gelation solution and incubated for 1.5 hours at 37°C under humidified conditions. Gels were transferred in denaturation buffer (PBS, 0.3 M Tris (VWR Chemicals, 28811.295); pH 8.8, 0.5 M NaCl (ThermoFisher Scientific), 14% SDS (Sigma-Aldrich, 05030-1L-F)) and autoclaved for 100 min. Next, the gels were washed in Milli-Q water (MQ) 3x 30 min to induce expansion (maximum up to 4x expansion), followed by 2x 15 min PBS washes, in preparation for antibody staining. All the washing steps were performed at room temperature.

#### Gel staining

##### Antibody staining

The gels were incubated in a blocking buffer (PBS, 2% bovine serum albumin (BSA) (ROTH, 8076.3), 2% normal goat serum (NGS) (abcam, ab7481), 1% normal donkey serum (NDS) (abcam, ab7475) for 3 hours at room temperature, with shaking. Next, samples were incubated in a primary antibody solution, prepared by diluting chicken anti-GFP antibody (Aves Lab, GFP-1010) at 1:250 ratio in antibody buffer (PBS, 2% BSA, 2% NGS, 1% NDS, 0.1% Triton X-100 (TX100) (Sigma-Aldrich, X100)), O/N at room temperature, with shaking. Gels were washed in washing buffer (PBS, 0.1% TX100) 3x 15 min at room temperature with shaking, and incubated for 3 hours at 37°C, with shaking with secondary goat anti-chicken Alexa Fluor 488 IgY (H+L) antibody (ThermoFisher Scientific, A-11039), diluted at 1:500 ratio in the antibody buffer. Gels were washed in the washing buffer 3x 15 min, followed 2× 10min PBS washes at room temperature, with shaking. Antibody staining, when applied, preceded pan staining.

##### Pan staining

Gels were incubated in 2 µg/ml ATTO 565 NHS-ester (ATTO-TEC GmbH, AD-565-31) for 2 hours, then washed in PBS, 2% BSA for 30 min, followed by 3× 10 min PBS washes. All the steps were performed at room temperature with shaking.

#### Sample mounting and microscopy

Gels were incubated in MQ 3x 30 min at room temperature, with shaking. The expanded gel was fixed at a poly-L-lysine-coated coverglass (VWR Avantor, 631-0172), placed in Attofluor imaging chamber (ThermoFisher, A7816). 1 ml of MQ was added to allow the gel to be completely covered.

The gel-samples were imaged using the LSM980 AiryScan 2 microscope (Zeiss), employing the SR-4Y Airy-Scan mode, and using the LD C-Apochromat 40x/1.1 water-immersion objective. Z-stacks were collected at a step size of 0.30 µm, at a typical pixel size of 62 nm.

The microscope was controlled by ZEN Blue v3.9 software and the images were subsequently processed with the super-resolution function in Auto mode with the SuperResolution factor of 4.70– 5.10 depending on the image. Scale bars and measured image dimensions were divided by the expansion factor to represent their corresponding pre-expansion sizes. 3D movies of expansion samples were rendered using BigTrace plugin for Fiji (Katrukha et al., 2025). To determine z-to-z distances, sarcomeres in single z planes were analyzed in Fiji. First, it was determined whether a sarcomere was in contracted or relaxed state. Next, the z-to-z distance was measured and values were divided by the expansion factor. Multiple z-to-z distances from one induvial were plotted distinguishing between contracted and relaxed states.

### Immuno-electron microscopy

#### Tissue collection

Adult zebrafish ventricles were isolated in extraction buffer (PBS, 10% KCL (BOOM, 7447-40-7), heparin (Sigma-Aldrich, 9041-08-1) and fixed in 4% PFA + 0.1% glutaraldehyde (Sigma-Aldrich, 111-30-8) in PHEM buffer (37 °C, 30 min). Subsequently, the hearts were washed 2× 10 min in NaBH_4_ (0.1 M) (Sigma-Aldrich, 16940-66-2) in PHEM buffer followed by 1x 30 min glycine (0.1 M) (ThermoFisher Scientific, 56-40-6) in PHEM buffer. Next, hearts were mounted in 1% agarose (Invitrogen, 16500-500) in 7×7×5 basemolds (VWR, 720-0820) and cardiac slices were made by using a vibratome (150 μm) (Leica VT 1200S). The cardiac slices were stored in 1% PFA in PHEM buffer at 4 °C until further use.

#### Sample preparation and immuno-electron microscopy

Cardiac slices were infiltrated with gelatin, cut into blocks and infused in 2.3 M sucrose O/N. The blocks were mounted on aluminum pins, snap-frozen in liquid nitrogen and sectioned at 70 nm using a UC7 cryo-ultramicrotome (Leica Microsystems). Sections were collected in a 1:1 mixture of methylcellulose and 2.3 M sucrose. For immunogold labelling, sections were washed in PBS at 37 °C for 30 min, followed by 3x 2 min incubations on drops of 0.15% glycine. Sections were then blocked for 5 min with 1% BSA-c/VG and incubated for 1 hour at room temperature with a primary antibody against Lamp1 (1:10, Boster Biological Technology, DZ01398-1). After washing, sections were incubated with Protein A conjugated to 10 nm gold particles (Cell Microscopy Core, UMC Utrecht). Finally, grids were contrasted with uranyl acetate and imaged in a Transmission Electron Microscope (Technai 12, Thermo Fischer Scientific) operating SerialEM software.

### Rapamycin treatment

Rapamycin (Sigma-Aldrich, 53123-88-9) treatments were performed as described previously (Chávez et al., 2020). In more detail, a 20 mg/ml stock solution was prepared in DMSO (Sigma-Aldrich, 67-68-5). Subsequently, a 0.3 mg/ml working solution was prepared in Cortland solution (Kinkel et al., 2010). Next, using a Hamilton syringe (gauge 30), a dose of 5 mg rapamycin (Sigma-Aldrich, 53123-88-9) was injected intraperitoneal per kg body weight. The fish received nine rapamycin (Sigma-Aldrich, 53123-88-9) doses which were injected every other day. Injections were performed as described previously (Kinkel et al., 2010). 2 hours after the last injection, zebrafish were euthanized and hearts were harvested and processed like described before.

### Western blot

Zebrafish ventricles were extracted in extraction buffer (PBS + heparin (Sigma-Aldrich, 9041-08-1)) and flash frozen in liquid nitrogen. A solution containing 30 μl RIPA (Thermo Scientific, 10017003), protease inhibitor (Sigma-Aldrich, 11873580001) and PhosSTOP (Sigma-Aldrich, 4906837001) was added to each ventricle followed by low frequency sonication (30 sec on, 30 sec off, 2-4 cycles (until dissolved), 4 °C) (Bioruptor Pico, Diagenode). Protein concentration was measured by PierceTM BCA protein assay kit (Thermofisher, A55864) following manufacturer’s instructions. 10 μg of protein (Sup figure 3) or 8 μg of protein (Sup figure 6) in combination with protein ladder (Thermofisher, 26619) were loaded into a pre-cast NuPageTM 4-12% BIS-Tris gel (Invitrogen, NP0336BOX) and separation was achieved at a constant voltage (160 V) for 1 h using NuPageTM MES SDS running buffer. Proteins were transferred to a nitrocellulose membrane (BioRad, 1704158) using Trans-Blot Turbo Transfer System (Biorad). Membrane was blocked in 5% milk dissolved in TBS-Tw (50 mM Tris-HCl pH 7.14,, 0.15 M NaCl, 0.1 % Tween 20) for 2 hours and after membrane was incubated O/N at 4°C and shaking with the primary antibody diluted in 3% BSA 1% TBS-Tween (LC3 (1:1000, NB100-2220), Tropomyosin (1:1000, Sigma-Aldrich, T9283). Next day, the membrane was washed three times with TBS-Tw buffer and incubated with secondary antibody diluted in blocking buffer (1:7500 RDye 800CW Goat anti-Rabbit IgG (92632211) and Goat anti-Mouse IgG (H+L) Alexa Fluor™ 680 (A21057) for 2 hours at room temperature. After washing 3x, membrane was imaged using Amersham™ Typhoon™ biomolecular imager. Data were analyzed by Fiji.

### Statistics and reproducibility

All data is represented as mean ± SD. All experiments were performed with at least three biological replicates per group. All analysis were performed blinded to genotype. Statistical analysis were performed using GraphPad Prism (GraphPad Software). Which statistical analysis was performed is indicated in the figure legends.

## Notes

### Competing Interest Statement

The authors have declared no competing interest.

## References

Abbas, M. T., Baba Ali, N., Farina, J. M., Mahmoud, A. K., Pereyra, M., Scalia, I. G., Kamel, M. A., Barry, T., Lester, S. J., Cannan, C. R., Mital, R., Wilansky, S., Freeman, W. K., Chao, C.-J., Alsidawi, S., Ayoub, C., & Arsanjani, R. (2024). Role of Genetics in Diagnosis and Management of Hypertrophic Cardiomyopathy: A Glimpse into the Future. Biomedicines, 12(3), 682. 10.3390/biomedicines12030682

Algül, S., Dorsch, L. M., Sorop, O., Vink, A., Michels, M., Dos Remedios, C. G., Dalinghaus, M., Merkus, D., Duncker, D. J., Kuster, D. W. D., & van der Velden, J. (2023). The microtubule signature in cardiac disease: Etiology, disease stage, and age dependency. Journal of Comparative Physiology. B, Biochemical, Systemic, and Environmental Physiology, 193(5), 581–595. 10.1007/s00360-023-01509-1

Arbelo, E., Protonotarios, A., Gimeno, J. R., Arbustini, E., Barriales-Villa, R., Basso, C., Bezzina, C. R., Biagini, E., Blom, N. A., de Boer, R. A., De Winter, T., Elliott, P. M., Flather, M., Garcia-Pavia, P., Haugaa, K. H., Ingles, J., Jurcut, R. O., Klaassen, S., Limongelli, G., … ESC Scientific Document Group. (2023). 2023 ESC Guidelines for the management of cardiomyopathies. European Heart Journal, 44(37), 3503–3626. 10.1093/eurheartj/ehad194

Arif, T., Qiu, J., Khademian, H., Lohithakshan, A., Menon, A., Menon, V., Slavinsky, M., Batignes, M., Lin, M., Sebra, R., Beaumont, K. G., Benson, D. L., Tzavaras, N., Ménager, M. M., & Ghaffari, S. (2025). Reversing lysosomal dysfunction restores youthful state in aged hematopoietic stem cells. Cell Stem Cell, 32(12), 1904–1922.e7. 10.1016/j.stem.2025.10.012

Austin, K. M., Trembley, M. A., Chandler, S. F., Sanders, S. P., Saffitz, J. E., Abrams, D. J., & Pu, W. T. (2019). Molecular mechanisms of arrhythmogenic cardiomyopathy. Nature reviews. Cardiology, 16(9), 519–537. 10.1038/s41569-019-0200-7

Baxi, K., Ghavidel, A., Waddell, B., Harkness, T. A., & de Carvalho, C. E. (2017). Regulation of Lysosomal Function by the DAF-16 Forkhead Transcription Factor Couples Reproduction to Aging in Caenorhabditis elegans. Genetics, 207(1), 83–101. 10.1534/genetics.117.204222

Berridge, M. J. (2003). Cardiac calcium signalling. Biochemical Society Transactions, 31(5), 930–933. 10.1042/bst0310930

Bers, D. M. (2002). Cardiac excitation–contraction coupling. Nature, 415(6868), 198–205. 10.1038/415198a

Bertozzi, A., Wu, C.-C., Nguyen, P. D., Vasudevarao, M. D., Mulaw, M. A., Koopman, C. D., de Boer, T. P., Bakkers, J., & Weidinger, G. (2021). Is zebrafish heart regeneration “complete”? Lineage-restricted cardiomyocytes proliferate to pre-injury numbers but some fail to differentiate in fibrotic hearts. Developmental Biology, 471, 106–118. 10.1016/j.ydbio.2020.12.004

Caporizzo, M. A., & Prosser, B. L. (2022). The microtubule cytoskeleton in cardiac mechanics and heart failure. Nature Reviews. Cardiology, 19(6), 364–378. 10.1038/s41569-022-00692-y

Chávez, M. N., Morales, R. A., López-Crisosto, C., Roa, J. C., Allende, M. L., & Lavandero, S. (2020). Autophagy Activation in Zebrafish Heart Regeneration. Scientific Reports, 10(1), 2191. 10.1038/s41598-020-59106-z

Chkourko, H. S., Guerrero-Serna, G., Lin, X., Darwish, N., Pohlmann, J. R., Cook, K. E., Martens, J. R., Rothenberg, E., Musa, H., & Delmar, M. (2012). Remodeling of mechanical junctions and of microtubule-associated proteins accompany cardiac connexin43 lateralization. Heart Rhythm, 9(7), 1133–1140.e6. 10.1016/j.hrthm.2012.03.003

Cleary, S. R., Teng, A. C. T., Kongmeneck, A. D., Fang, X., Phillips, T. A., Cho, E. E., Smith, R. A., Karkut, P., Makarewich, C. A., Kekenes-Huskey, P. M., Gramolini, A. O., & Robia, S. L. (2025). Dilated cardiomyopathy variant R14del increases phospholamban pentamer stability, blunting dynamic regulation of calcium. The Journal of Biological Chemistry, 301(2), 108118. 10.1016/j.jbc.2024.108118

Colacurcio, D. J., & Nixon, R. A. (2016). Disorders of lysosomal acidification—The emerging role of v-ATPase in aging and neurodegenerative disease. Ageing Research Reviews, Lysosomes in Aging, 32, 75–88. 10.1016/j.arr.2016.05.004

Costa, S., Cerrone, M., Saguner, A. M., Brunckhorst, C., Delmar, M., & Duru, F. (2021). Arrhythmogenic cardiomyopathy: An in-depth look at molecular mechanisms and clinical correlates. Trends in Cardiovascular Medicine, 31(7), 395–402. 10.1016/j.tcm.2020.07.006

Damstra, H. G., Mohar, B., Eddison, M., Akhmanova, A., Kapitein, L. C., & Tillberg, P. W. (2022). Visualizing cellular and tissue ultrastructure using Ten-fold Robust Expansion Microscopy (TREx). eLife, 11, e73775. 10.7554/eLife.73775

Dees, E., Miller, P. M., Moynihan, K. L., Pooley, R. D., Hunt, R. P., Galindo, C. L., Rottman, J. N., & Bader, D. M. (2012). Cardiac-specific deletion of the microtubule-binding protein CENP-F causes dilated cardiomyopathy. Disease Models & Mechanisms, 5(4), 468–480. 10.1242/dmm.008680

Delmar, M., & McKenna, W. J. (2010). The cardiac desmosome and arrhythmogenic cardiomyopathies: From gene to disease. Circulation Research, 107(6), 700–714. 10.1161/CIRCRESAHA.110.223412

Dutzmann, J., Kefalianakis, Z., Kahles, F., Daniel, J.-M., Gufler, H., Wohlgemuth, W. A., Knöpp, K., & Sedding, D. G. (2024). Intermittent Fasting After ST-Segment–Elevation Myocardial Infarction Improves Left Ventricular Function: The Randomized Controlled INTERFAST-MI Trial. Circulation: Heart Failure, 17(5), e010936. 10.1161/CIRCHEARTFAILURE.123.010936

Eijgenraam, T. R., Boogerd, C. J., Stege, N. M., Oliveira Nunes Teixeira, V., Dokter, M. M., Schmidt, L. E., Yin, X., Theofilatos, K., Mayr, M., van der Meer, P., van Rooij, E., van der Velden, J., Silljé, H. H. W., & de Boer, R. A. (2021). Protein Aggregation Is an Early Manifestation of Phospholamban p.(Arg14del)–Related Cardiomyopathy: Development of PLN-R14del–Related Cardiomyopathy. Circulation: Heart Failure, 14(11), e008532. 10.1161/CIRCHEARTFAILURE.121.008532

Fedeli, C., Filadi, R., Rossi, A., Mammucari, C., & Pizzo, P. (2019). PSEN2 (presenilin 2) mutants linked to familial Alzheimer disease impair autophagy by altering Ca2+ homeostasis. Autophagy, 15(12), 2044–2062. 10.1080/15548627.2019.1596489

Felice, T. (2017). Pompe Disease, a Storage Cardiomyopathy. Cardiogenetics, 7(1), 6857. 10.4081/cardiogenetics.2017.6857

Gambarotto, Davide, Hamel, Virginie, & Guichard, Paul. (2021). Ultrastructure expansion microscopy (U-ExM). In Methods in Cell Biology (Vol. 161, pp. 57–81). Elsevier. 10.1016/bs.mcb.2020.05.006

Gerull, B., & Brodehl, A. (2020). Genetic Animal Models for Arrhythmogenic Cardiomyopathy. Frontiers in Physiology, 11. 10.3389/fphys.2020.00624

Gilbert, G., Demydenko, K., Dries, E., Puertas, R. D., Jin, X., Sipido, K., & Roderick, H. L. (2020). Calcium Signaling in Cardiomyocyte Function. Cold Spring Harbor Perspectives in Biology, 12(3), a035428. 10.1101/cshperspect.a035428

Godar, R. J., Ma, X., Liu, H., Murphy, J. T., Weinheimer, C. J., Kovacs, A., Crosby, S. D., Saftig, P., & Diwan, A. (2015). Repetitive stimulation of autophagy-lysosome machinery by intermittent fasting preconditions the myocardium to ischemia-reperfusion injury. Autophagy, 11(9), 1537–1560. 10.1080/15548627.2015.1063768

Haghighi, K., Gardner, G., Vafiadaki, E., Kumar, M., Green, L. C., Ma, J., Crocker, J. S., Koch, S., Arvanitis, D. A., Bidwell, P., Rubinstein, J., van de Leur, R., Doevendans, P. A., Akar, F. G., Tranter, M., Wang, H.-S., Sadayappan, S., DeMazumder, D., Sanoudou, D., … Kranias, E. G. (2021). Impaired Right Ventricular Calcium Cycling Is an Early Risk Factor in R14del-Phospholamban Arrhythmias. Journal of Personalized Medicine, 11(6), 502. 10.3390/jpm11060502

Haghighi, K., Kolokathis, F., Gramolini, A. O., Waggoner, J. R., Pater, L., Lynch, R. A., Fan, G.-C., Tsiapras, D., Parekh, R. R., Dorn, G. W., MacLennan, D. H., Kremastinos, D. T., & Kranias, E. G. (2006). A mutation in the human phospholamban gene, deleting arginine 14, results in lethal, hereditary cardiomyopathy. Proceedings of the National Academy of Sciences of the United States of America, 103(5), 1388–1393. 10.1073/pnas.0510519103

Hayashi, Y., Takatori, S., Warsame, W. Y., Tomita, T., Fujisawa, T., & Ichijo, H. (2023). TOLLIP acts as a cargo adaptor to promote lysosomal degradation of aberrant ER membrane proteins. The EMBO Journal, 42(23), e114272. 10.15252/embj.2023114272

Hof, I. E., van der Heijden, J. F., Kranias, E. G., Sanoudou, D., de Boer, R. A., van Tintelen, J. P., van der Zwaag, P. A., & Doevendans, P. A. (2019). Prevalence and cardiac phenotype of patients with a phospholamban mutation. Netherlands Heart Journal: Monthly Journal of the Netherlands Society of Cardiology and the Netherlands Heart Foundation, 27(2), 64–69. 10.1007/s12471-018-1211-4

Honkoop, H., de Bakker, D. E., Aharonov, A., Kruse, F., Shakked, A., Nguyen, P. D., de Heus, C., Garric, L., Muraro, M. J., Shoffner, A., Tessadori, F., Peterson, J. C., Noort, W., Bertozzi, A., Weidinger, G., Posthuma, G., Grün, D., van der Laarse, W. J., Klumperman, J., … Bakkers, J. (2019). Single-cell analysis uncovers that metabolic reprogramming by ErbB2 signaling is essential for cardiomyocyte proliferation in the regenerating heart. eLife, 8, e50163. 10.7554/eLife.50163

Honkoop, H., Nguyen, P. D., van der Velden, V. E. M., Sonnen, K. F., & Bakkers, J. (2021). Live imaging of adult zebrafish cardiomyocyte proliferation ex vivo. Development (Cambridge, England), 148(18), dev199740. 10.1242/dev.199740

Jung, S., Chung, Y., Lee, Y., Lee, Y., Cho, J. W., Shin, E.-J., Kim, H.-C., & Oh, Y. J. (2019). Buffering of cytosolic calcium plays a neuroprotective role by preserving the autophagy-lysosome pathway during MPP+-induced neuronal death. Cell Death Discovery, 5, 130. 10.1038/s41420-019-0210-6

Kamel, S. M., van Opbergen, C. J. M., Koopman, C. D., Verkerk, A. O., Boukens, B. J. D., de Jonge, B., Onderwater, Y. L., van Alebeek, E., Chocron, S., Polidoro Pontalti, C., Weuring, W. J., Vos, M. A., de Boer, T. P., van Veen, T. A. B., & Bakkers, J. (2021). Istaroxime treatment ameliorates calcium dysregulation in a zebrafish model of phospholamban R14del cardiomyopathy. Nature Communications, 12, 7151. 10.1038/s41467-021-27461-8

Katrukha, E., Gros, A., & Eglinger, J. (2025). ekatrukha/BigTrace: V.0.7.0 [Software]. Zenodo. 10.5281/zenodo.17185904

Kemmler, C. L., Moran, H. R., Murray, B. F., Scoresby, A., Klem, J. R., Eckert, R. L., Lepovsky, E., Bertho, S., Nieuwenhuize, S., Burger, S., D’Agati, G., Betz, C., Puller, A.-C., Felker, A., Ditrychova, K., Bötschi, S., Affolter, M., Rohner, N., Lovely, C. B., … Mosimann, C. (2023). Next-generation plasmids for transgenesis in zebrafish and beyond. Development, 150(8), dev201531. 10.1242/dev.201531

Kim, Y., Lee, Y., Choo, M., Yun, N., Cho, J. W., & Oh, Y. J. (2023). A surge of cytosolic calcium dysregulates lysosomal function and impairs autophagy flux during cupric chloride–induced neuronal death. The Journal of Biological Chemistry, 300(1), 105479. 10.1016/j.jbc.2023.105479

Kinkel, M. D., Eames, S. C., Philipson, L. H., & Prince, V. E. (2010). Intraperitoneal Injection into Adult Zebrafish. Journal of Visualized Experiments (JoVE), (42), e2126. 10.3791/2126

Kumar, M., Haghighi, K., Koch, S., Rubinstein, J., Stillitano, F., Hajjar, R. J., Kranias, E. G., & Sadayappan, S. (2023). Myofilament Alterations Associated with Human R14del-Phospholamban Cardiomyopathy. International Journal of Molecular Sciences, 24(3), 2675. 10.3390/ijms24032675

Kwan, K. M., Fujimoto, E., Grabher, C., Mangum, B. D., Hardy, M. E., Campbell, D. S., Parant, J. M., Yost, H. J., Kanki, J. P., & Chien, C.-B. (2007). The Tol2kit: A multisite gateway-based construction kit for Tol2 transposon transgenesis constructs. Developmental Dynamics, 236(11), 3088–3099. 10.1002/dvdy.21343

Lazzarini, E., Jongbloed, J. D. H., Pilichou, K., Thiene, G., Basso, C., Bikker, H., Charbon, B., Swertz, M., van Tintelen, J. P., & van der Zwaag, P. A. (2015). The ARVD/C Genetic Variants Database: 2014 Update. Human Mutation, 36(4), 403–410. 10.1002/humu.22765

Lee, A. S. (2005). The ER chaperone and signaling regulator GRP78/BiP as a monitor of endoplasmic reticulum stress. Methods, 35(4), 373–381. 10.1016/j.ymeth.2004.10.010

Lim, S. H. Y., Hansen, M., & Kumsta, C. (2024). Molecular Mechanisms of Autophagy Decline during Aging. Cells, 13(16), 1364. 10.3390/cells13161364

Lin, Y.-F., Swinburne, I., & Yelon, D. (2012). Multiple influences of blood flow on cardiomyocyte hypertrophy in the embryonic zebrafish heart. Developmental Biology, 362(2), 242–253. 10.1016/j.ydbio.2011.12.005

Ma, X., Mani, K., Liu, H., Kovacs, A., Murphy, J. T., Foroughi, L., French, B. A., Weinheimer, C. J., Kraja, A., Benjamin, I. J., Hill, J. A., Javaheri, A., & Diwan, A. (2019). Transcription Factor EB Activation Rescues Advanced αB-Crystallin Mutation-Induced Cardiomyopathy by Normalizing Desmin Localization. Journal of the American Heart Association: Cardiovascular and Cerebrovascular Disease, 8(4), e010866. 10.1161/JAHA.118.010866

Macquart, C., Jüttner, R., Morales Rodriguez, B., Le Dour, C., Lefebvre, F., Chatzifrangkeskou, M., Schmitt, A., Gotthardt, M., Bonne, G., & Muchir, A. (2019). Microtubule cytoskeleton regulates Connexin 43 localization and cardiac conduction in cardiomyopathy caused by mutation in A-type lamins gene. Human Molecular Genetics, 28(24), 4043–4052. 10.1093/hmg/ddy227

Maniezzi, C., Eskandr, M., Florindi, C., Ferrandi, M., Barassi, P., Sacco, E., Pasquale, V., Maione, A. S., Pompilio, G., Teixeira, V. O. N., de Boer, R. A., Silljé, H. H. W., Lodola, F., & Zaza, A. (2024). Early consequences of the phospholamban mutation PLN-R14del+/− in a transgenic mouse model. Acta Physiologica (Oxford, England), 240(3), e14082. 10.1111/apha.14082

Marwaha, R., & Sharma, M. (2017). DQ-Red BSA Trafficking Assay in Cultured Cells to Assess Cargo Delivery to Lysosomes. Bio-Protocol, 7(19), e2571. 10.21769/BioProtoc.2571

McGuire, C. M., & Forgac, M. (2018). Glucose starvation increases V-ATPase assembly and activity in mammalian cells through AMP kinase and phosphatidylinositide 3-kinase/Akt signaling. Journal of Biological Chemistry, 293(23), 9113–9123. 10.1074/jbc.RA117.001327

McNally, E. M., & Mestroni, L. (2017). Dilated Cardiomyopathy. Circulation Research, 121(7), 731–748. 10.1161/CIRCRESAHA.116.309396

McNally, E., MacLeod, H., & Dellefave-Castillo, L. (1993). Arrhythmogenic Right Ventricular Cardiomyopathy Overview. In M. P. Adam, S. Bick, G. M. Mirzaa, R. A. Pagon, S. E. Wallace, & A. Amemiya (Red.), GeneReviews®. University of Washington, Seattle. http://www.ncbi.nlm.nih.gov/books/NBK1131/

Muraro, M. J., Dharmadhikari, G., Grün, D., Groen, N., Dielen, T., Jansen, E., van Gurp, L., Engelse, M. A., Carlotti, F., de Koning, E. J. P., & van Oudenaarden, A. (2016). A Single-Cell Transcriptome Atlas of the Human Pancreas. Cell Systems, 3(4), 385–394.e3. 10.1016/j.cels.2016.09.002

Mustaly-Kalimi, S., Gallegos, W., Marr, R. A., Gilman-Sachs, A., Peterson, D. A., Sekler, I., & Stutzmann, G. E. (2022). Protein mishandling and impaired lysosomal proteolysis generated through calcium dysregulation in Alzheimer’s disease. Proceedings of the National Academy of Sciences, 119(49), e2211999119. 10.1073/pnas.2211999119

Nguyen, P. D., Gooijers, I., Campostrini, G., Verkerk, A. O., Honkoop, H., Bouwman, M., de Bakker, D. E. M., Koopmans, T., Vink, A., Lamers, G. E. M., Shakked, A., Mars, J., Mulder, A. A., Chocron, S., Bartscherer, K., Tzahor, E., Mummery, C. L., de Boer, T. P., Bellin, M., & Bakkers, J. (2023). Interplay between calcium and sarcomeres directs cardiomyocyte maturation during regeneration. Science, 380(6646), 758–764. 10.1126/science.abo6718

Nixon, R. A. (2020). The aging lysosome: An essential catalyst for late-onset neurodegenerative diseases. Biochimica Et Biophysica Acta. Proteins and Proteomics, 1868(9), 140443. 10.1016/j.bbapap.2020.140443

Olzmann, J. A., Li, L., & Chin, L. S. (2008). Aggresome formation and neurodegenerative diseases: Therapeutic implications. Current Medicinal Chemistry, 15(1), 47–60. 10.2174/092986708783330692

Pan, B., Zhang, H., Cui, T., & Wang, X. (2017). TFEB activation protects against cardiac proteotoxicity via increasing autophagic flux. Journal of Molecular and Cellular Cardiology, 113, 51–62. 10.1016/j.yjmcc.2017.10.003

Patel, D. M., Dubash, A. D., Kreitzer, G., & Green, K. J. (2014). Disease mutations in desmoplakin inhibit Cx43 membrane targeting mediated by desmoplakin-EB1 interactions. The Journal of Cell Biology, 206(6), 779–797. 10.1083/jcb.201312110

Ragone, I., Barallobre-Barreiro, J., Takov, K., Theofilatos, K., Yin, X., Schmidt, L. E., Domenech, N., Crespo-Leiro, M. G., van der Voorn, S. M., Vink, A., van Veen, T. A. B., Bödör, C., Merkely, B., Radovits, T., & Mayr, M. (2023). SERCA2a Protein Levels Are Unaltered in Human Heart Failure. Circulation, 148(7), 613–616. 10.1161/CIRCULATIONAHA.123.064513

Ramdas, N. M., & Shivashankar, G. V. (2015). Cytoskeletal Control of Nuclear Morphology and Chromatin Organization. Journal of Molecular Biology, Functional Relevance and Dynamics of Nuclear Organization, 427(3), 695–706. 10.1016/j.jmb.2014.09.008

Salomo-Coll, C., Jimenez-Moreno, N., & Wilkinson, S. (2025). Lysosomal Degradation of ER Client Proteins by ER-phagy and Related Pathways. Journal of Molecular Biology, 169035. 10.1016/j.jmb.2025.169035

Sasazawa, Y., Date, Y., Hattori, N., & Saiki, S. (2024). Clustering lysosomes around the MTOC: A promising strategy for SNCA/alpha-synuclein breakdown leading to parkinson disease treatment. Autophagy, 20(12), 2839–2840. 10.1080/15548627.2024.2413295

Schmitt, J. P., Kamisago, M., Asahi, M., Li, G. H., Ahmad, F., Mende, U., Kranias, E. G., MacLennan, D. H., Seidman, J. G., & Seidman, C. E. (2003). Dilated Cardiomyopathy and Heart Failure Caused by a Mutation in Phospholamban. Science, 299(5611), 1410–1413. 10.1126/science.1081578

Sciarretta, S., Yee, D., Nagarajan, N., Bianchi, F., Saito, T., Valenti, V., Tong, M., Del Re, D. P., Vecchione, C., Schirone, L., Forte, M., Rubattu, S., Shirakabe, A., Boppana, V. S., Volpe, M., Frati, G., Zhai, P., & Sadoshima, J. (2018). Trehalose-Induced Activation of Autophagy Improves Cardiac Remodeling After Myocardial Infarction. Journal of the American College of Cardiology, 71(18), 1999–2010. 10.1016/j.jacc.2018.02.066

Stege, N. M., de Boer, R. A., Makarewich, C. A., van der Meer, P., & Silljé, H. H. W. (2024). Reassessing the Mechanisms of PLN-R14del Cardiomyopathy: From Calcium Dysregulation to S/ER Malformation. JACC: Basic to Translational Science, 9(8), 1041–1052. 10.1016/j.jacbts.2024.02.017

Stege, N. M., Eijgenraam, T. R., Oliveira Nunes Teixeira, V., Feringa, A. M., Schouten, E. M., Kuster, D. W. D., van der Velden, J., Wolters, A. H. G., Giepmans, B. N. G., Makarewich, C. A., Bassel-Duby, R., Olson, E. N., de Boer, R. A., & Silljé, H. H. W. (2023). DWORF Extends Life Span in a PLN-R14del Cardiomyopathy Mouse Model by Reducing Abnormal Sarcoplasmic Reticulum Clusters. Circulation Research, 133(12), 1006–1021. 10.1161/CIRCRESAHA.123.323304

Stypmann, J., Gläser, K., Roth, W., Tobin, D. J., Petermann, I., Matthias, R., Mönnig, G., Haverkamp, W., Breithardt, G., Schmahl, W., Peters, C., & Reinheckel, T. (2002). Dilated cardiomyopathy in mice deficient for the lysosomal cysteine peptidase cathepsin L. Proceedings of the National Academy of Sciences of the United States of America, 99(9), 6234–6239. 10.1073/pnas.092637699

Tan, R., Foster, P. J., Needleman, D. J., & McKenney, R. J. (2018). Cooperative Accumulation of Dynein-Dynactin at Microtubule Minus-Ends Drives Microtubule Network Reorganization. Developmental Cell, 44(2), 233–247.e4. 10.1016/j.devcel.2017.12.023

Tantama, M., Martínez-François, J. R., Mongeon, R., & Yellen, G. (2013). Imaging energy status in live cells with a fluorescent biosensor of the intracellular ATP-to-ADP ratio. Nature Communications, 4, 2550. 10.1038/ncomms3550

Te Rijdt, W. P., Asimaki, A., Jongbloed, J. D. H., Hoorntje, E. T., Lazzarini, E., van der Zwaag, P. A., de Boer, R. A., van Tintelen, J. P., Saffitz, J. E., van den Berg, M. P., & Suurmeijer, A. J. H. (2019). Distinct molecular signature of phospholamban p.Arg14del arrhythmogenic cardiomyopathy. Cardiovascular Pathology: The Official Journal of the Society for Cardiovascular Pathology, 40, 2–6. 10.1016/j.carpath.2018.12.006

te Rijdt, W. P., van der Klooster, Z. J., Hoorntje, E. T., Jongbloed, J. D. H., van der Zwaag, P. A., Asselbergs, F. W., Dooijes, D., de Boer, R. A., van Tintelen, J. P., van den Berg, M. P., Vink, A., & Suurmeijer, A. J. H. (2017). Phospholamban immunostaining is a highly sensitive and specific method for diagnosing phospholamban p.Arg14del cardiomyopathy. Cardiovascular Pathology, 30, 23–26. 10.1016/j.carpath.2017.05.004

te Rijdt, W. P., van Tintelen, J. P., Vink, A., van der Wal, A. C., de Boer, R. A., van den Berg, M. P., & Suurmeijer, A. J. H. (2016). Phospholamban p.Arg14del cardiomyopathy is characterized by phospholamban aggregates, aggresomes, and autophagic degradation. Histopathology, 69(4), 542–550. 10.1111/his.12963

Tessadori, F., Roessler, H. I., Savelberg, S. M. C., Chocron, S., Kamel, S. M., Duran, K. J., van Haelst, M. M., van Haaften, G., & Bakkers, J. (2018). Effective CRISPR/Cas9-based nucleotide editing in zebrafish to model human genetic cardiovascular disorders. Disease Models & Mechanisms, 11(10), dmm035469. 10.1242/dmm.035469

Triolo, M., & Hood, D. A. (2021). Manifestations of Age on Autophagy, Mitophagy and Lysosomes in Skeletal Muscle. Cells, 10(5), 1054. 10.3390/cells10051054

Uchida, K., Scarborough, E. A., & Prosser, B. L. (2022). Cardiomyocyte Microtubules: Control of Mechanics, Transport, and Remodeling. Annual review of physiology, 84, 257–283. 10.1146/annurev-physiol-062421-040656

Vafiadaki, E., Glijnis, P. C., Doevendans, P. A., Kranias, E. G., & Sanoudou, D. (2023). Phospholamban R14del disease: The past, the present and the future. Frontiers in Cardiovascular Medicine, 10. 10.3389/fcvm.2023.1162205

Vafiadaki, E., Haghighi, K., Arvanitis, D. A., Kranias, E. G., & Sanoudou, D. (2022). Aberrant PLN-R14del Protein Interactions Intensify SERCA2a Inhibition, Driving Impaired Ca2+ Handling and Arrhythmogenesis. International Journal of Molecular Sciences, 23(13), 6947. 10.3390/ijms23136947

Vafiadaki, E., Kranias, E. G., Eliopoulos, A. G., & Sanoudou, D. (2024). The phospholamban R14del generates pathogenic aggregates by impairing autophagosome-lysosome fusion. Cellular and Molecular Life Sciences: CMLS, 81(1), 450. 10.1007/s00018-024-05471-1

van der Beek, J., de Heus, C., Sanza, P., Liv, N., & Klumperman, J. (2024). Loss of the HOPS complex disrupts early-to-late endosome transition, impairs endosomal recycling and induces accumulation of amphisomes. Molecular Biology of the Cell, 35(3), ar40. 10.1091/mbc.E23-08-0328

van der Zwaag, P. A., van Rijsingen, I. A. W., Asimaki, A., Jongbloed, J. D. H., van Veldhuisen, D. J., Wiesfeld, A. C. P., Cox, M. G. P. J., van Lochem, L. T., de Boer, R. A., Hofstra, R. M. W., Christiaans, I., van Spaendonck-Zwarts, K. Y., Lekanne dit Deprez, R. H., Judge, D. P., Calkins, H., Suurmeijer, A. J. H., Hauer, R. N. W., Saffitz, J. E., Wilde, A. A. M., … van Tintelen, J. P. (2012). Phospholamban R14del mutation in patients diagnosed with dilated cardiomyopathy or arrhythmogenic right ventricular cardiomyopathy: Evidence supporting the concept of arrhythmogenic cardiomyopathy. European Journal of Heart Failure, 14(11), 1199–1207. 10.1093/eurjhf/hfs119

van der Zwaag, P. A., van Rijsingen, I. A. W., de Ruiter, R., Nannenberg, E. A., Groeneweg, J. A., Post, J. G., Hauer, R. N. W., van Gelder, I. C., van den Berg, M. P., van der Harst, P., Wilde, A. A. M., & van Tintelen, J. P. (2013). Recurrent and founder mutations in the Netherlands— Phospholamban p.Arg14del mutation causes arrhythmogenic cardiomyopathy. Netherlands Heart Journal, 21(6), 286–293. 10.1007/s12471-013-0401-3

van Opbergen, C. J. M., Koopman, C. D., Kok, B. J. M., Knöpfel, T., Renninger, S. L., Orger, M. B., Vos, M. A., van Veen, T. A. B., Bakkers, J., & de Boer, T. P. (2018). Optogenetic sensors in the zebrafish heart: A novel in vivo electrophysiological tool to study cardiac arrhythmogenesis. Theranostics, 8(17), 4750–4764. 10.7150/thno.26108

van Rijsingen, I. A. W., van der Zwaag, P. A., Groeneweg, J. A., Nannenberg, E. A., Jongbloed, J. D. H., Zwinderman, A. H., Pinto, Y. M., dit Deprez, R. H. L., Post, J. G., Tan, H. L., de Boer, R. A., Hauer, R. N. W., Christiaans, I., van den Berg, M. P., van Tintelen, J. P., & Wilde, A. A. M. (2014). Outcome in Phospholamban R14del Carriers. Circulation: Cardiovascular Genetics, 7(4), 455–465. 10.1161/CIRCGENETICS.113.000374

Wu, A. Z., Xu, D., Yang, N., Lin, S.-F., Chen, P.-S., Cala, S. E., & Chen, Z. (2016). Phospholamban is concentrated in the nuclear envelope of cardiomyocytes and involved in perinuclear/nuclear calcium handling. Journal of Molecular and Cellular Cardiology, 100, 1–8. 10.1016/j.yjmcc.2016.09.008

Wu, L., Zhao, Y., Gong, X., Liang, Z., Yu, J., Wang, J., Zhang, Y., Wang, X., Shu, X., & Bao, J. (2025). Intermittent Fasting Ameliorates β-Amyloid Deposition and Cognitive Impairment Accompanied by Decreased Lipid Droplet Aggregation Within Microglia in an Alzheimer’s Disease Model. Molecular Nutrition & Food Research, 69(4), e202400660. 10.1002/mnfr.202400660

Yang, Z., McMahon, C. J., Smith, L. R., Bersola, J., Adesina, A. M., Breinholt, J. P., Kearney, D. L., Dreyer, W. J., Denfield, S. W., Price, J. F., Grenier, M., Kertesz, N. J., Clunie, S. K., Fernbach, S. D., Southern, J. F., Berger, S., Towbin, J. A., Bowles, K. R., & Bowles, N. E. (2005). Danon Disease as an Underrecognized Cause of Hypertrophic Cardiomyopathy in Children. Circulation, 112(11), 1612–1617. 10.1161/CIRCULATIONAHA.105.546481

Young, J. E., Martinez, R. A., & La Spada, A. R. (2009). Nutrient Deprivation Induces Neuronal Autophagy and Implicates Reduced Insulin Signaling in Neuroprotective Autophagy Activation*. Journal of Biological Chemistry, 284(4), 2363–2373. 10.1074/jbc.M806088200

