## Supplemental Figures for "Fasting reverses PLN R14del-mediated cardiomyopathy through lysosomal reactivation"

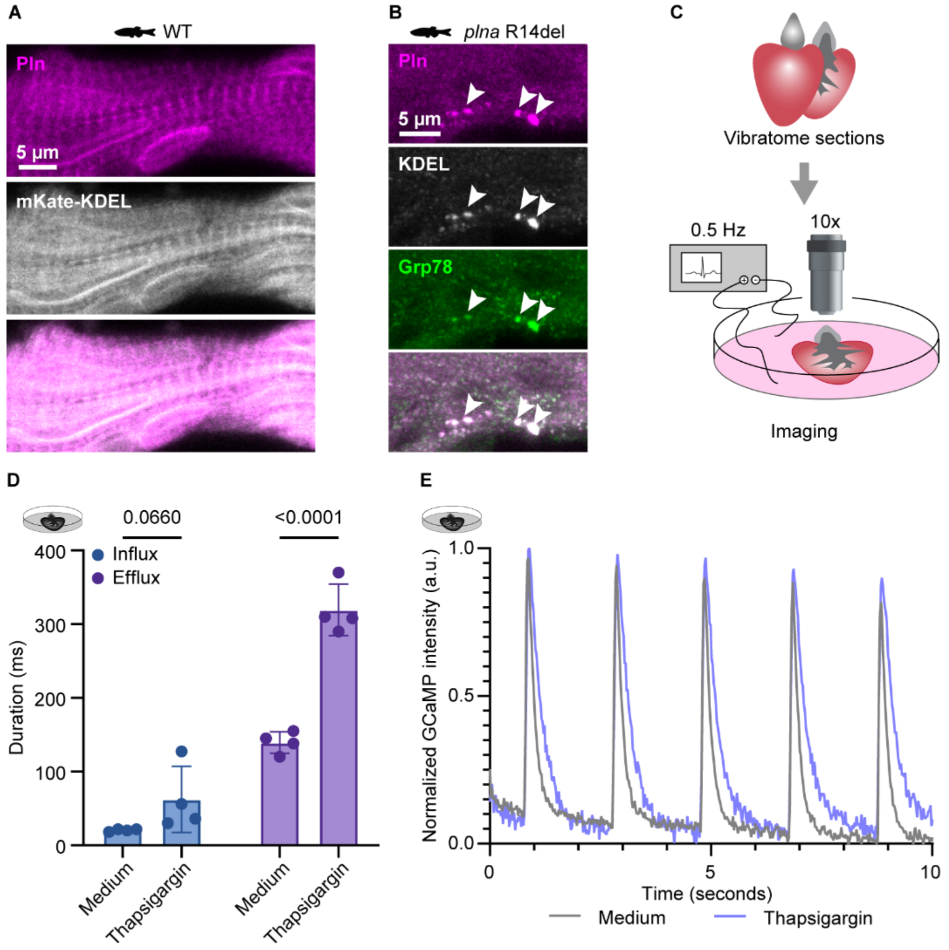


**Supplemental figure 1. mKate-KDEL localization, SR stress evaluation in *plna* R14del hearts and validation of thapsigargin treatment. A.** Representative images of immunofluorescent staining against Pln and mKate-KDEL in one year old wild type (WT) Tg(*myl7*:mKate-KDEL) hearts. Data shown are representative for n = 3 WT hearts. Scale bar = 5 µm. **B.** Representative images of immunofluorescent staining against Pln, mKate-KDEL and Grp78 in one year old *plna* R14del Tg(*myl7*:mKate-KDEL;exorh:GFP) zebrafish hearts. Data shown are representative for n = 3 *plna* R14del hearts. Scale bar = 5 µm. **C.** Schematic overview of the paced cardiac slices; Tg(*myl7*:GCaMP6f-nls-T2A-RCaMP107-NES-pA)$^{hu11799}$ zebrafish hearts were extracted and sectioned into 150 µm thick slices using a vibratome. The slices were then cultured and attached to electrical stimulation (0.5 Hz). Subsequently, the GCaMP signal was recorded at highspeed *ex vivo*. **D.** Quantification of $Ca^{2+}$ influx and $Ca^{2+}$ efflux in WT cardiac slices treated with medium or thapsigargin (100 µM). Data points represent individual hearts with n = 4 hearts per group. Error bars indicate mean ± s.d. Statistics were performed by two-way ANOVA. **E.** Representative images $Ca^{2+}$ peaks of a WT slices treated with medium (grey) and thapsigargin (100 µM) (blue). Data shown are representative for n = 4 hearts per group.

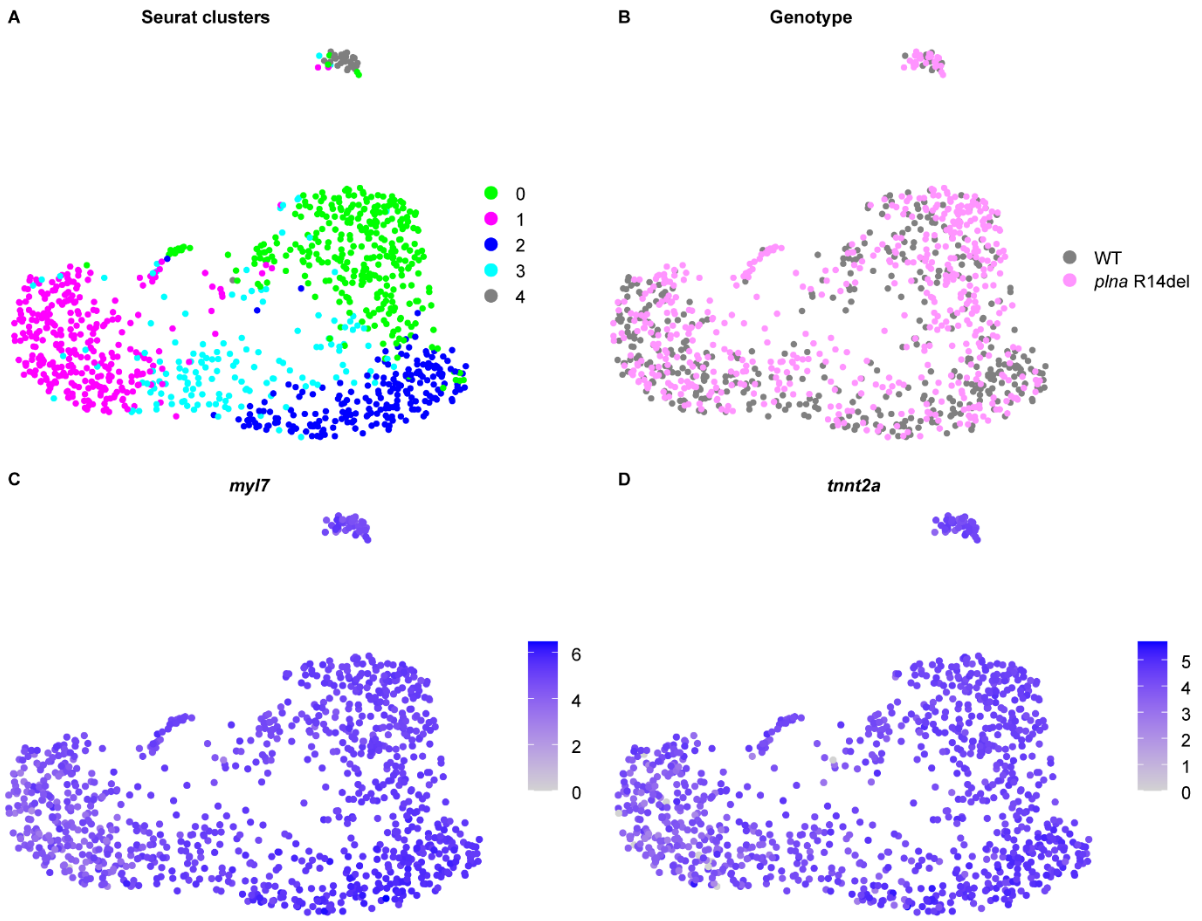


E

| Clusters | 0 | 1 | 2 | 3 | 4 |
|---|---|---|---|---|---|
| WT | 38.5% | 51.2% | 67.1% | 47.5% | 45.5% |
| *plna* R14del | 61.5% | 48.8% | 32.9% | 52.5% | 54.5% |

**Supplemental figure 2. UMAP of SORT-seq dataset showing a main cardiomyocyte cluster. A.** Uniform Manifold Approximation and Projection (UMAP) representation of sorted cells with high cpmVenus fluorescence from one year old wild type (WT) and *plna* R14del Tg(*myl7*:PercevalHR) hearts. **B.** UMAP representation of transcriptome similarities between individual WT (grey) and *plna* R14del (magenta) zebrafish cardiomyocytes from one year old fish. **C-D.** UMAP representation of expression of the cardiac genes *myl7* **(C)** and *tnnt2a* **(D)** in sorted cells. **E.** Normalized percentages of WT and *plna* R14del cells per cluster, calculated by weighting both the relative abundance of each genotype within the cluster and the total number of cells per genotype. This approach accounts for differences in total cell numbers between WT and *plna* R14del cells, allowing direct comparison of their distributions across clusters.

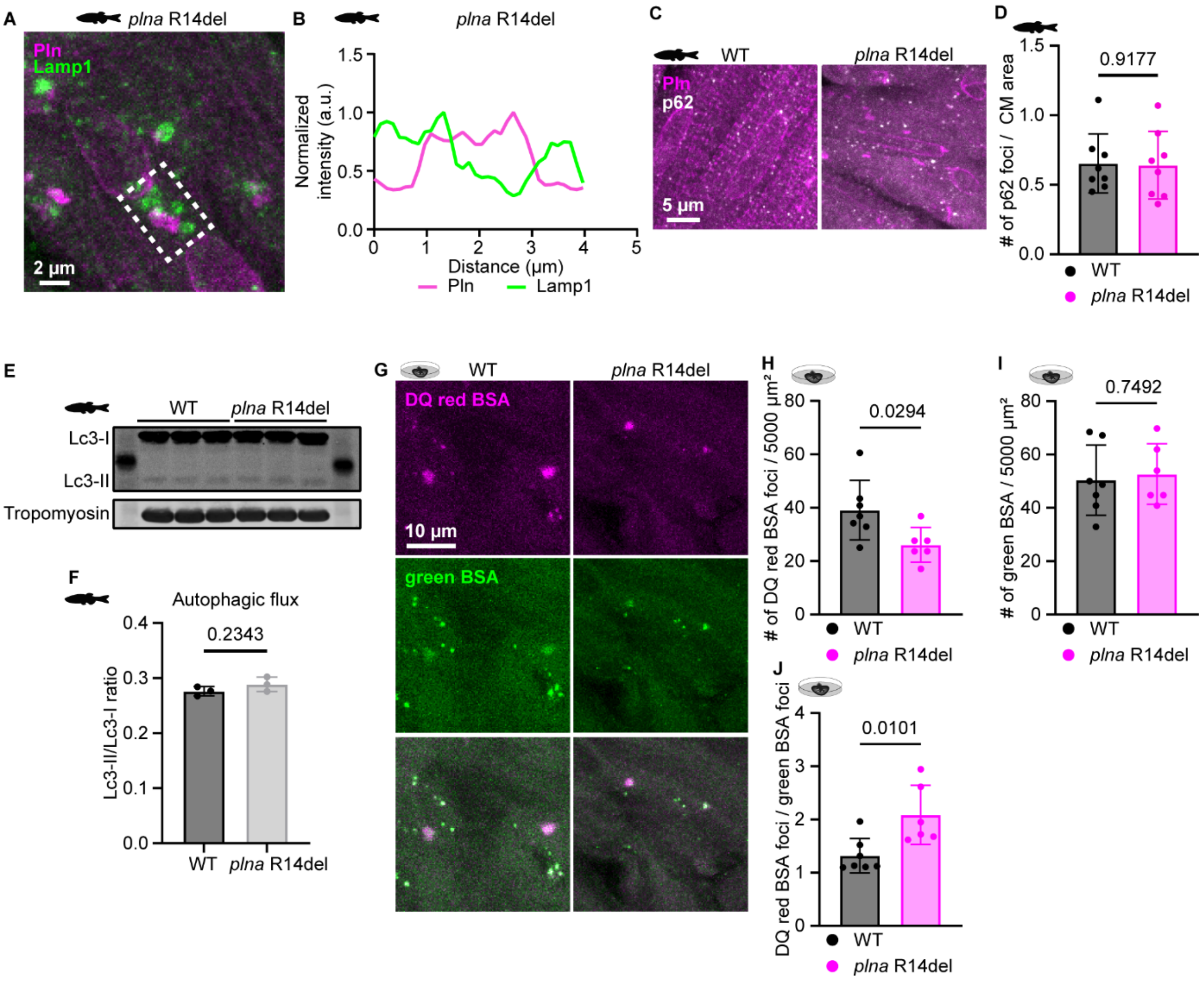


**Supplemental figure 3. Pln localization near Lamp1 foci in *plna* R14del hearts and evaluation of macro-autophagy and endocytosis in WT and *plna* R14del hearts. A.** Representative images of immunofluorescent staining against Pln and Lamp1 in one year old *plna* R14del zebrafish. Dashed white box indicates region used for plot line in panel B. Data shown is representative for n = 8 *plna* R14del zebrafish hearts. Scale bar = 2 µm. **B.** Representative line plots showing Pln (magenta) and Lamp1 (green) adjacent co-localization in *plna* R14del cardiomyocytes. **C.** Representative images of immunofluorescent staining against Pln and p62 in one year old wild type (WT) and *plna* R14del hearts. p62 marks autophagosomes. Data shown are representative for n = 8 WT and n = 8 *plna* R14del zebrafish hearts. Scale bar = 5 µm. **D**. Quantification of number of p62 foci divided by cardiomyocyte (CM) area. Data points represent individual hearts with n = 8 per group. Error bars indicate mean ± s.d. Statistics were performed by two-tailed unpaired t-test. **E.** Western blot against Lc3-I, Lc3-II and Tropomyosin in one year old WT and *plna* R14del zebrafish hearts. n = 3 biological replicates were included per group. The ladder indicates a band at 15 kDa. **F.** Quantification of Lc3-II intensity divided by Lc3-I intensity from the western blot representative for the autophagy flux in one year old WT and *plna* R14del zebrafish hearts. Data points represent individual hearts with n = 3 hearts included per group. Error bars indicate mean ± s.d. Statistics were performed by two-tailed unpaired t-test. **G.** Representative images of DQ red BSA and green BSA *ex vivo* assay in one year old WT and *plna* R14del zebrafish hearts. DQ red BSA dots represent functional lysosomes while green BSA dots represent all endocytic vesicles while. Data shown are representative for n = 3 WT hearts and n = 3 *plna* R14del hearts. Scale bar = 10 µm. **H.** Quantification of number of DQ red BSA foci per 5000 µm$^2$. Data points represent an individual measurement per image; multiple points originate from one individual; n = 3 individuals were included in both groups. Error bars indicate mean ± s.d. Statistics were performed by two-tailed unpaired t-test. **I.** Quantification of number of green BSA foci per 5000 µm$^2$. Data points represent an individual measurement per image; multiple points originate from one individual; n = 3 individuals were included in both groups. Error bars indicate mean ± s.d. Statistics were performed by two-tailed unpaired t-test. **J.** Quantification of DQ red BSA foci / green BSA foci. Data points represent an individual measurement per image; multiple points originate from one individual; n = 3 individuals were included in both groups. Error bars indicate mean ± s.d. Statistics were performed by two-tailed unpaired t-test.

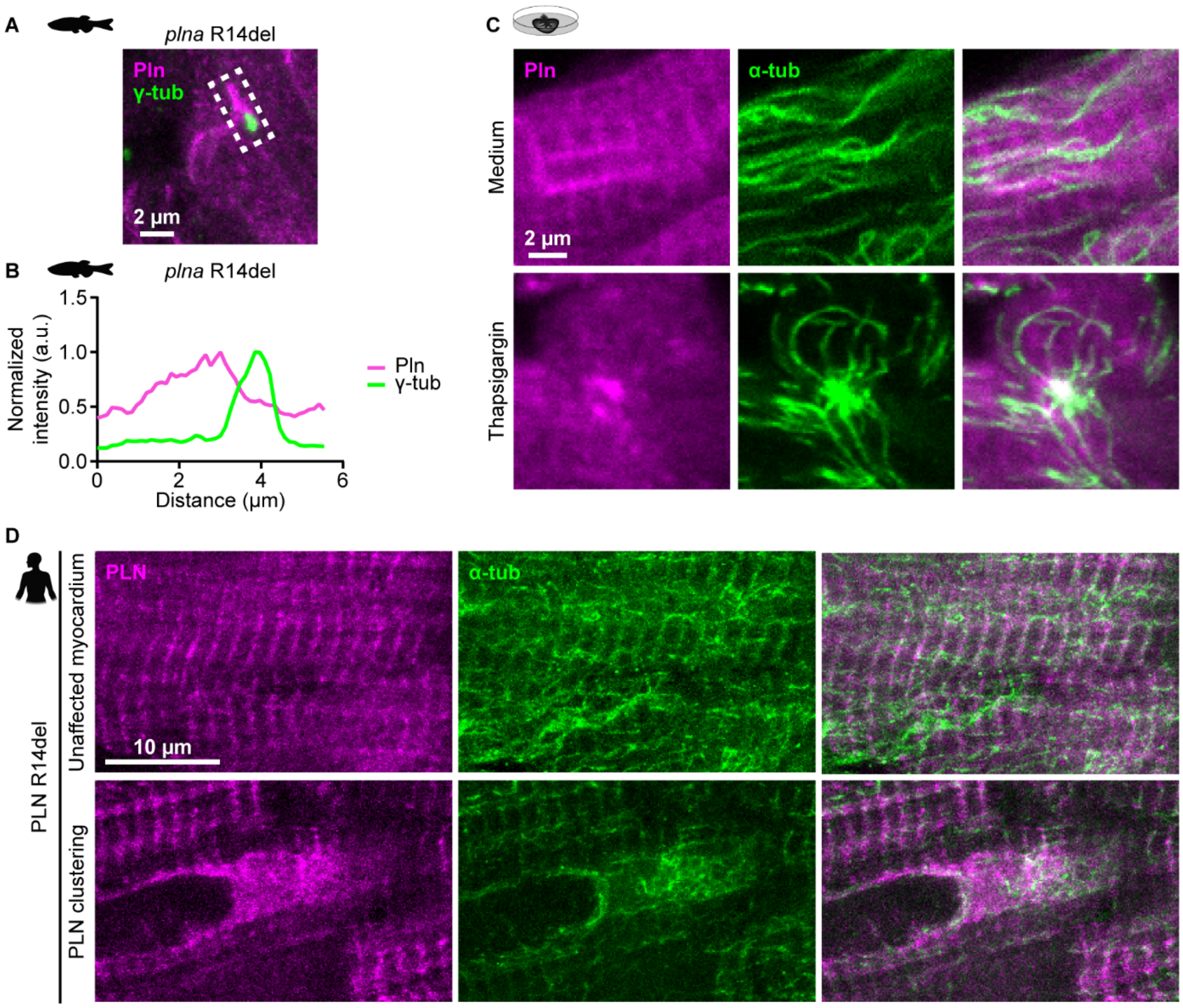


**Supplemental figure 4. Pln localization at the MTOC in *plna* R14del hearts and microtubules reorganization in WT zebrafish cardiac slices treated with thapsigargin as well as human PLN R14del patients. A.** Representative images of immunofluorescent staining against Pln and γ-tubulin (γ-tub) in one year old *plna* R14del zebrafish. Dashed white box indicates region used for plot line in panel B. Scale bar = 2 µm. **B.** Representative line plots showing Pln (magenta) and γ-tubulin (green) adjacent co-localization in *plna* R14del cardiomyocytes. **C.** Representative images of immunofluorescent staining against Pln and α-tubulin (α-tub) in wild type (WT) zebrafish cardiac slices treated with medium or thapsigargin (100 µM). Data shown are representative for n = 3 hearts per group. Scale bar = 2 µm. **D.** Representative images of immunofluorescent staining against PLN and α-tub in left ventricular (LV) tissue obtained from patients with end-stage cardiomyopathy carrying the PLN R14del variant. Top panels show unaffected myocardium, while below panels show a region containing PLN clustering. Data shown are representative for n = 4 individuals. Scale bar = 10 µm.

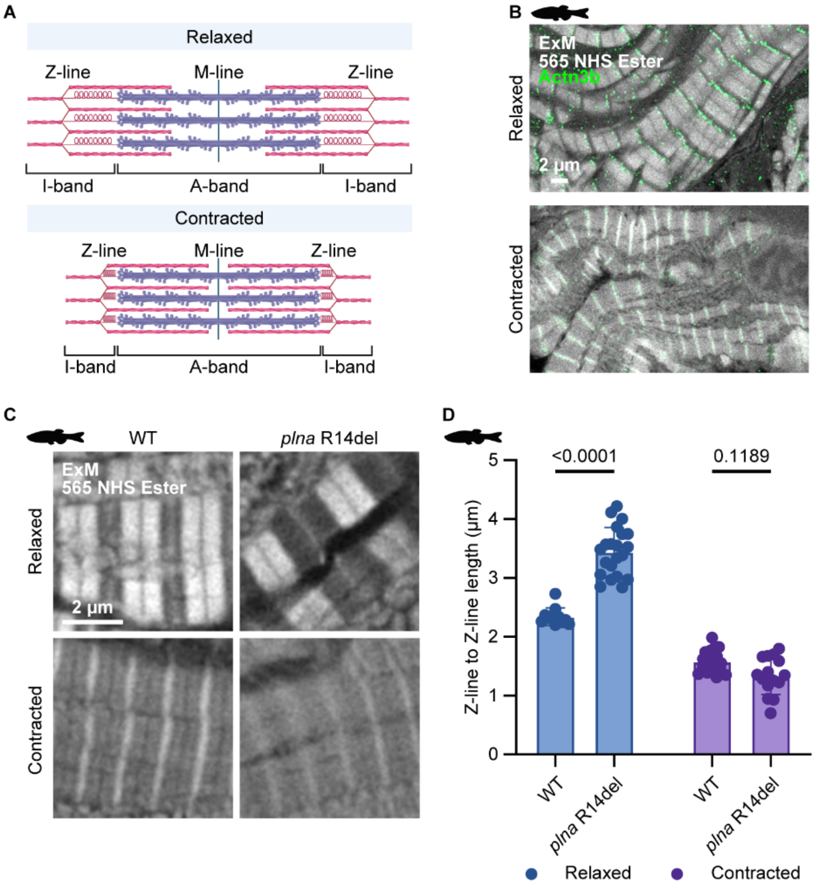


**Supplemental figure 5. ExM allows for detailed evaluation of sarcomere structure and shows that *plna* R14del cardiomyocytes have enlarged Z-line to Z-line space in the relaxed state. A.** Schematic overview of relaxed and contracted sarcomeres. Z-line, M-line, I-band and A-band are indicated. **B.** Representative images of 565 NHS ester general protein stain in Tg(*myl7:actn3b*-EGFP) expanded zebrafish hearts. Actn3b is indicated in green and localizes to the Z-line. Data shown are representative for n = 2 zebrafish hearts. Scale bar = 2 µm; scale bar is corrected for pre-expansion dimensions. **C.** Representative images of relaxed and contracted sarcomeres visualized by the 565 NHS ester general protein stain in one year old wild type (WT) and *plna* R14del expanded zebrafish hearts. Data shown are representative for n = 3 WT and n = 3 *plna* R14del zebrafish hearts. Scale bar = 2 µm; scale bar is corrected for pre-expansion dimensions. **D.** Quantification of Z-line to Z-line length in µm in one year old WT and *plna* R14del zebrafish in the relaxed and contracted state. The measured length is corrected for pre-expansion dimensions. Data points represent an individual measurement; all points originate from one individual. Measurements were performed twice in two different individuals, yielding similar results. Error bars indicate mean ± s.d. Statistics were performed by two-way ANOVA.

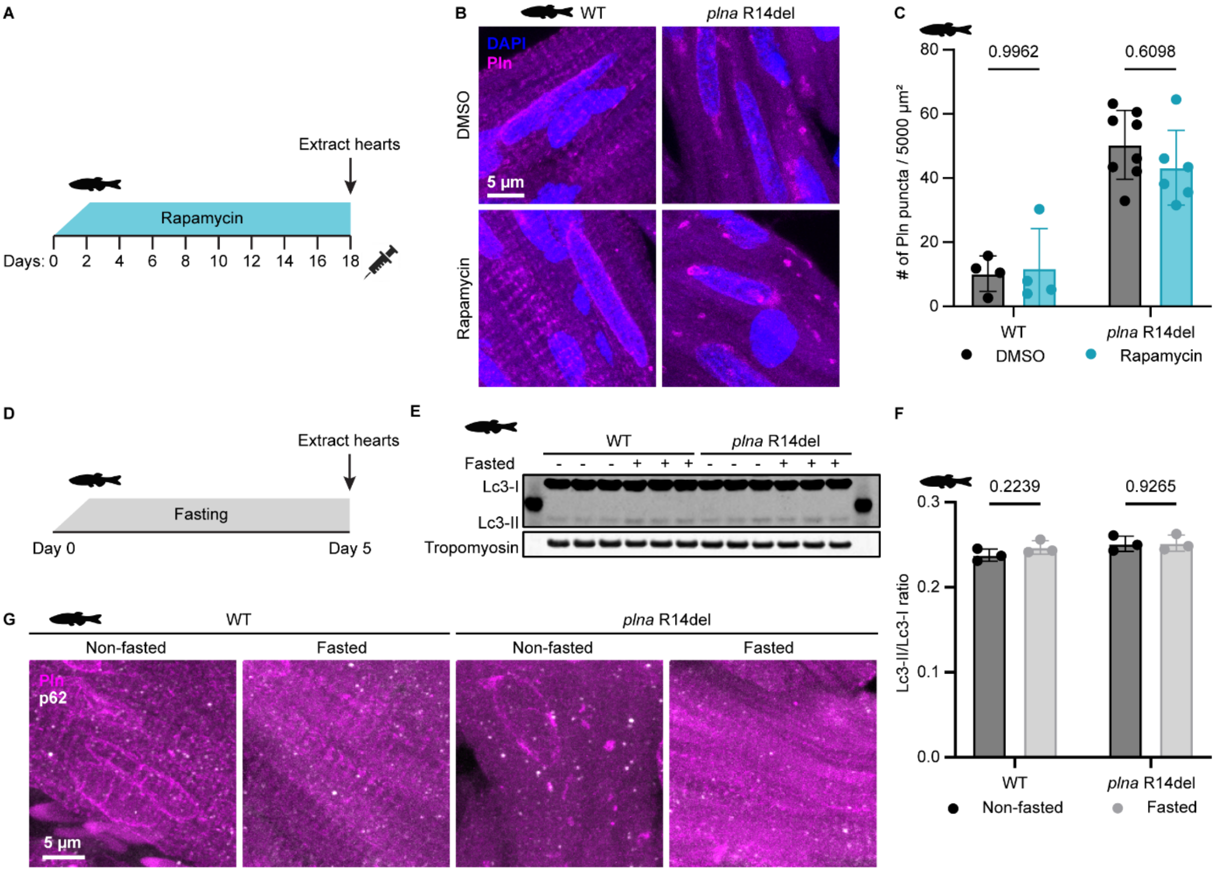


**Supplemental figure 6. Rescue of SR-derived vesicle accumulation upon fasting is independent of macro-autophagy. A.** Workflow rapamycin 5 mg/kg treatment in wild type (WT) and *plna* R14del one year old zebrafish. **B.** Representative images of immunofluorescent staining against DAPI and Pln in one year old DMSO and rapamycin treated WT and *plna* R14del zebrafish. Data shown are representative for at least n = 4 individuals per group. Scale bar = 5 µm. **C.** Quantification of Pln puncta per 5000 µm$^2$ in one year old DMSO or rapamycin treated WT and *plna* R14del zebrafish. Data points represent individual hearts with at least n = 4 per group. Error bars indicate mean ± s.d. Statistics were performed by two-way ANOVA. **D.** Workflow fasting in WT and *plna* R14del one year old zebrafish. **E.** Western blot against Lc3-I, Lc3-II and Tropomyosin in one year old non-fasted versus fasted WT and *plna* R14del zebrafish hearts. n = 3 biological replicates were included per group. The ladder indicates a band at 15 kDa. **F.** Quantification of Lc3-II intensity divided by Lc3-I intensity from the western blot representative for the autophagy flux in one year old non-fasted versus fasted WT and *plna* R14del zebrafish hearts. Data points represent individual hearts with n = 3 hearts included per group. Error bars indicate mean ± s.d. Statistics were performed by two-way ANOVA. **G.** Representative images of immunofluorescent staining against Pln and p62 in one year old non-fasted versus fasted WT and *plna* R14del hearts. p62 marks autophagosomes. Data shown are representative for at least n = 6 individuals per group. Scale bar = 5 µm.

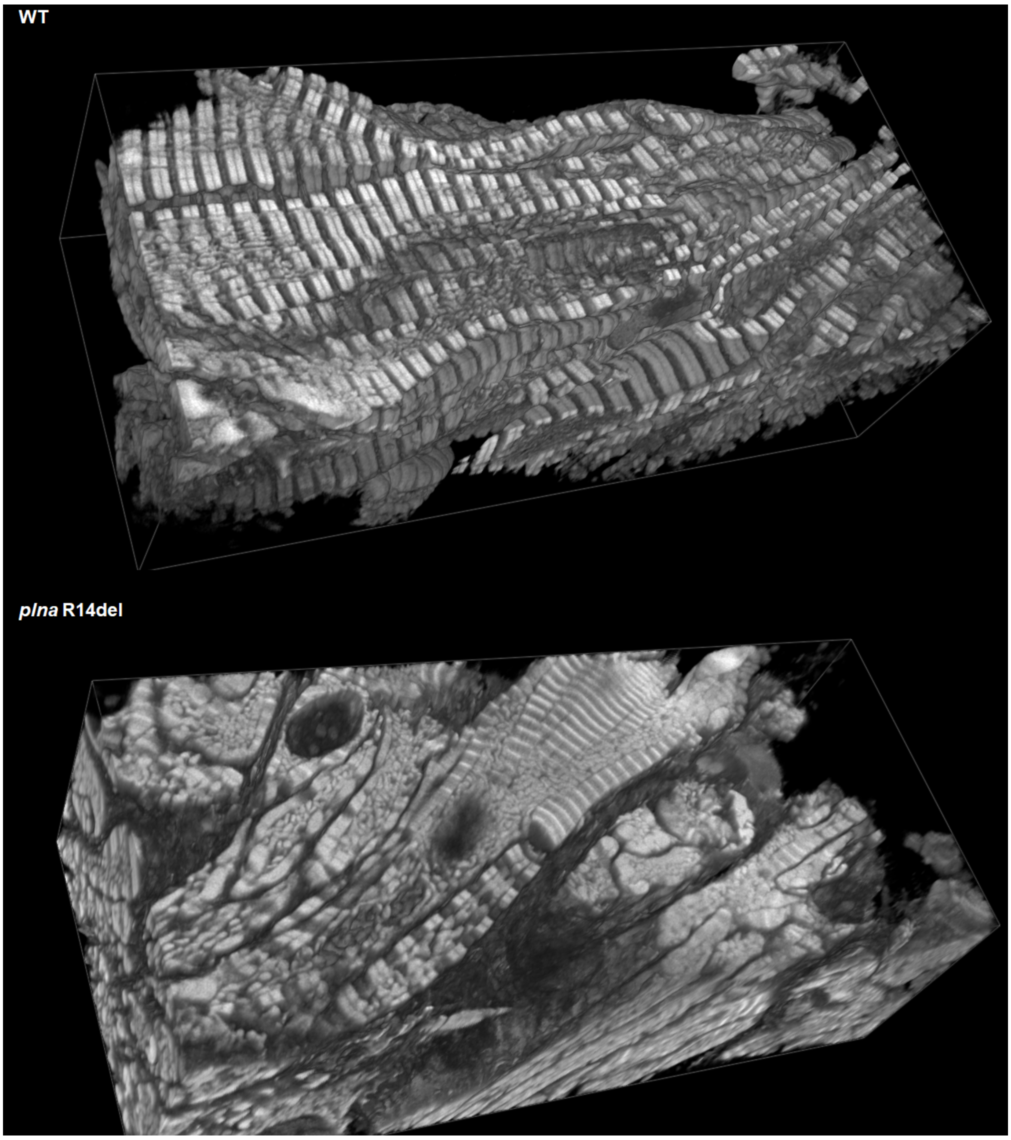


**Supplemental video 1. 3D visualization of expanded WT and *plna* R14del zebrafish cardiac slices.** 3D videos of 565 NHS ester general protein stain in one year old wild type (WT) and *plna* R14del expanded zebrafish hearts shown in figure 4A. Pre-expansion corrected dimensions of shown regions are 73 × 32 × 23 µm (X × Y × Z). 3D videos were rendered using BigTrace plugin for Fiji (Katrukha et al., 2025).
